# Scanning ion conductance microscopy reveals differences in the ionic environments of gram positive and negative bacteria

**DOI:** 10.1101/2020.08.26.267849

**Authors:** Kelsey Cremin, Bryn Jones, James Teahan, Gabriel N. Meloni, David Perry, Christian Zerfass, Munehiro Asally, Orkun S. Soyer, Patrick R. Unwin

## Abstract

This paper reports on the use of scanning ion conductance microscopy (SICM) to locally map the ionic properties and charge environment of two live bacterial strains: the gramnegative *Escherichia coli* and the gram-positive *Bacillus subtilis*. SICM results find heterogeneities across the bacterial surface, and significant differences among the grampositive and -negative bacteria. The bioelectrical environment of the *B. subtilis* was found to be considerably more negatively charged compared to *E. coli*. SICM measurements, fitted to a simplified finite element method (FEM) model, revealed surface charge values of −80 to −140 mC m^−2^ for the gram-negative *E. coli*. The gram-positive *B. subtilis* show a much higher conductivity around the cell wall, and surface charge values between −350 and −450 mC m^−2^ were found using the same simplified model. SICM was also able to detect regions of high negative charge near *B. subtilis*, not detected in the topographical SICM response and attributed to extracellular polymeric substance. To further explore how the *B. subtilis* cell wall structure can influence the SICM current response, a more comprehensive FEM model, accounting for the physical properties of the gram-positive cell wall, was developed. The new model provides a more realistic description of the cell wall and allowed investigation of the relation between its key properties and SICM currents, building foundations to further investigate and improve understanding of the gram-positive cellular microenvironment.

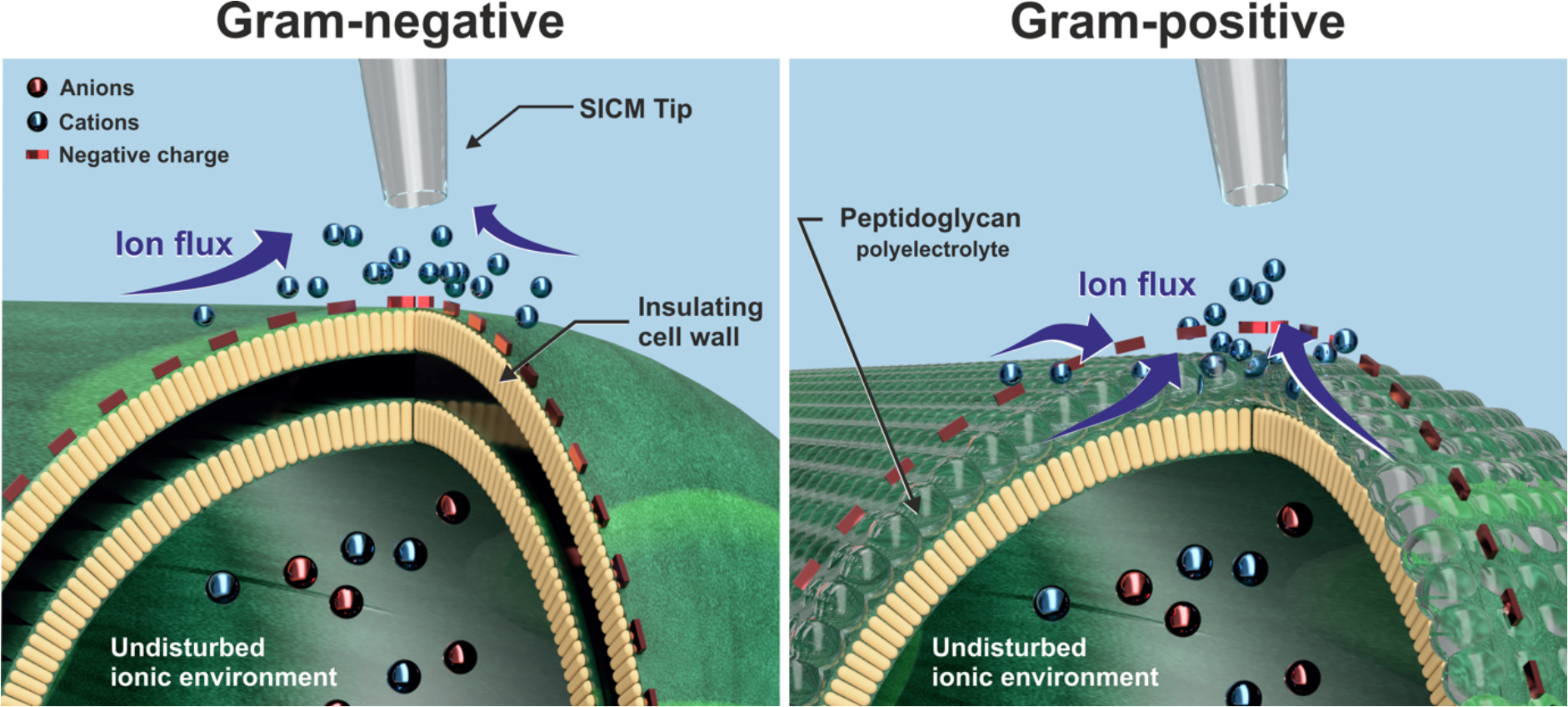

## Introduction

Scanning ion conductance microscopy (SICM) is proving to be increasingly versatile for nanoscale functional mapping of live biological entities *in situ* to provide an abundance of information about the surface topography,^1,2^ together with interfacial properties,^3^ cell junction permeability,^4–7^ and dynamic processes.^8–12^ SICM utilizes a glass or quartz nanopipette, filled with electrolytic solution, to probe a substrate immersed in an electrolyte bath. A bias is applied between two quasi-reference counter electrodes (QRCEs); one in the nanopipette and the other in the bulk solution to drive an ionic current through the nanopipette. The current magnitude is highly sensitive to nanopipette-substrate separation and local ion conductivity, including that arising from the double layer at charged interfaces, and can be used to probe (independently) topography and surface charge of a range of substrates.^13–15^

Critical information about the ionic environment of the substrate -solution interface can be acquired by employing specific potential-time SICM scanning protocols,^16,17^ in tandem with finite element method (FEM) simulations, as exemplified by the quantitative analysis of surface charge^15,16,18–20^ and for reaction mapping.^12,21^ In this paper, we consider the use of SICM to characterize the ionic (bioelectrical) environment of single live bacterial cells. There is increasing interest in characterizing the bacterial cell microenvironment,^22^ as membrane properties and extracellular charge distributions are considered to play a role in biofilm formation,^23^ nutrient uptake,^24^ and cell differentiation (e.g. persister cells and biofilm forming cells).^25,26^

The bacterial microenvironment is expected to be directly influenced by the bacterial cell wall through surface appendages and proteins, as well as cellular secretions. The cell wall of bacteria can be broadly classified into two polyphyletic groups; gram-positive and gramnegative (Figure 1), each arising from structural differences which underpin the bacterial interactions with the environment.^27^ As seen in Figure 1, gram-negative cell walls have an additional membrane covered in lipopolysaccharides, while gram-positive walls contain only one membrane and have a thicker layer of peptidoglycan containing negatively charged teichoic acids.^28^

**Figure 1:**
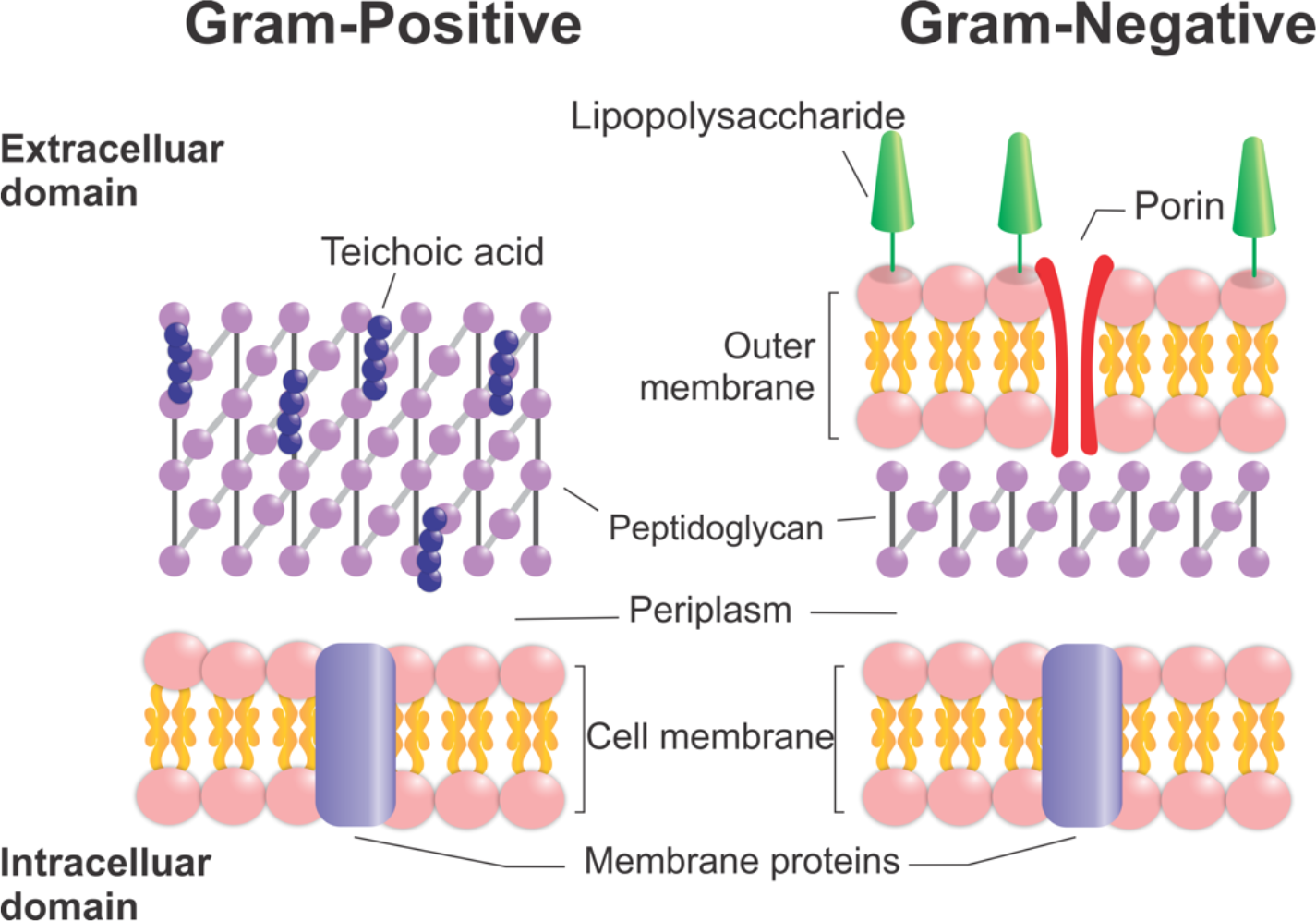
Cartoon illustration of bacterial cell envelope highlighting its key components in gram-positive and gram-negative bacteria.

Previous zeta-potential and electrophoretic measurements have shown that grampositive and gram-negative bacteria both have a net negative surface charge, but the magnitude of the charge density is different between these two broad groups, and between individual species within the groups.^29–31^ Zeta-potential measurements determine the electric charge at the shear plane of the solid-liquid interface, and do not provide a holistic descriptor of the charge environment at the cell wall, including important factors such as ionic permeability. Furthermore, the zeta-potential is measured for a population of cells, and is blind to possible heterogeneities between different cells or across the cell surface of a single bacterium. Atomic force microscopy (AFM) can be applied to surface charge mapping of single cells,^32^ as done for microbial cells.^33^ AFM is only sensitive to the charge environment of the outermost surface, whereas with SICM there is the possibility of probing ion permeability of a sample.^4,34^ Furthermore, for non-invasive scanning of soft biological samples, SICM avoids deformation of the sample surface and consequently can provide better spatial resolution and longer viable experiment time scale.^35,36^

Here, we use SICM to map the charge environment of two different bacterial species, the gram-negative *Escherchia coli* and gram-positive *Bacillus subtilis*. Concurrent to these measurements, we develop a model of the properties and mechanisms influencing the interfacial charge and ion fluxes at and around the bacterial cell wall and perform FEM simulations to analyze model behavior against experimental data. Our overarching aims are to demonstrate the application of SICM to single cells of live bacteria, so to better quantify single cell microenvironment in gram-positive and gram-negative bacteria, and to provide a foundation that will enable future analyzes that can link diverse cellular behaviors to ionic dynamics within the cell-microenvironment interface.

## Experimental Methods

### Chemicals

All reagents were of analytical grade and used without further purification. Deionized water (Milli-Q, resistivity ca. 18.2 MΩ.cm at 25 °C) was used in the preparation of all solutions. 50 mM potassium chloride, buffered at pH 7.0 with tris(hydroxymethyl)aminomethane (both from Sigma-Aldrich) was used as supporting electrolyte for all the SICM experiments. Low melting point agarose (Cleaver Scientific, CSL-LMA100), poly-L-lysine (Sigma Aldrich) or Cell-Tak (Corning) were used to adhere bacteria to glass slides, with the use of each defined herein.

### Bacterial cultures

*B. subtilis* (NCIB 3610 - *Δhag* depleted (created by Daniel Kearns and obtained from Munehiro Asally, University of Warwick),^37^ and *E. coli K12* (wild type, obtained from DSMZ (DSM No. 498)) were cultured in a modified M9 media containing 0.4 % w/v glucose - full strain and media composition can be found in the *Supporting Information* (SI), Section SI-1 and 2. Bacteria were taken from freezer stocks (50 % glycerol, −80 ^o^C) and grown in 40 mL volumes of media, in sterile Erlenmeyer flasks. The cultures were grown overnight prior to SICM measurements on a shaking incubator at 37 ^o^C and 150 rpm.

### Bacterial substrates

Different adhesion methods were able to anchor and restrict inherent bacterial movement, whilst not inhibiting culture survival, as detailed in Section SI-3 and SI-4. In brief, adhesive layers of poly-L-lysine (PLL), Cell-Tak, or < 0.5 mm agarose gels were deposited on the glass surface of a 50 mm glass bottomed dish (WillCo Wells, USA, HBST-5040). A 100 μL aliquot of an overnight culture (optical density at 600 nm ~ 0.45) was drop cast to the adhesive substrate of choice. The sample was then left for 30 minutes at room temperature to adhere, followed by one gentle 10 mL application of clean media, which was then withdrawn by Pasteur pipette to remove any unadhered bacteria.

### SICM scanning regimes

Nanopipettes utilized as the scanning probes for SICM were fabricated using standard protocols and characterized by electron microscopy (EM). Fabrication and characterization protocols are described in Section SI-5. The end lumen radius of the nanopipettes was typically in the range 85-100 nm (measured accurately by EM). For SICM mapping, described in detail elsewhere,^15,16,18,19^ the nanopipette current was recorded continuously, while the nanopipette position and/or potential was varied synchronously. The nanopipette was approached towards the substrate/bacterial cell surface at a predefined position with a small applied bias between the internal and bulk QRCEs (*V_Tip_*, Figure 2A), typically 50 mV, where the ionic current response is primarily sensitive to nanopipette-substrate separation (Figure 2B, period I).^38^ Upon a 2 % decrease in current magnitude compared to the bulk value, corresponding to a nanopipette-substrate separation of tens of nm, as determined from FEM simulations (further explained in SI-6), the nanopipette movement stopped automatically, with the corresponding *z*-position (at each *x-y* position) revealing the substrate topography. With the nanopipette at this position, *V_Tip_* was either pulsed to −500 mV for a defined period; or scanned linearly to −500 mV, reversed linearly to 500 mV, and then back to the approach potential of 50 mV (Figure 2B, period II) before retracting the nanopipette by 4 μm, back into the bulk solution. The same potential pulsed or cyclic linear sweep program was repeated in the bulk (Figure 2B III and IV) to allow each surface electrochemical measurement to be normalized against a bulk measurement at each position (position-level self-referencing).^16,39^ Normalized current data are presented herein, where the near-surface current (at approx. 100 nm from substrate) values were divided by their corresponding bulk current (at approx. 4 μm from substrate) values. For the pulsed-potential regime, this normalization was done using data from the end of the respective pulsed potential period, whilst for the potential scans each datapoint in the current-potential plot at the surface was normalized by the equivalent scan datapoint obtained in bulk solution.

**Figure 2:**
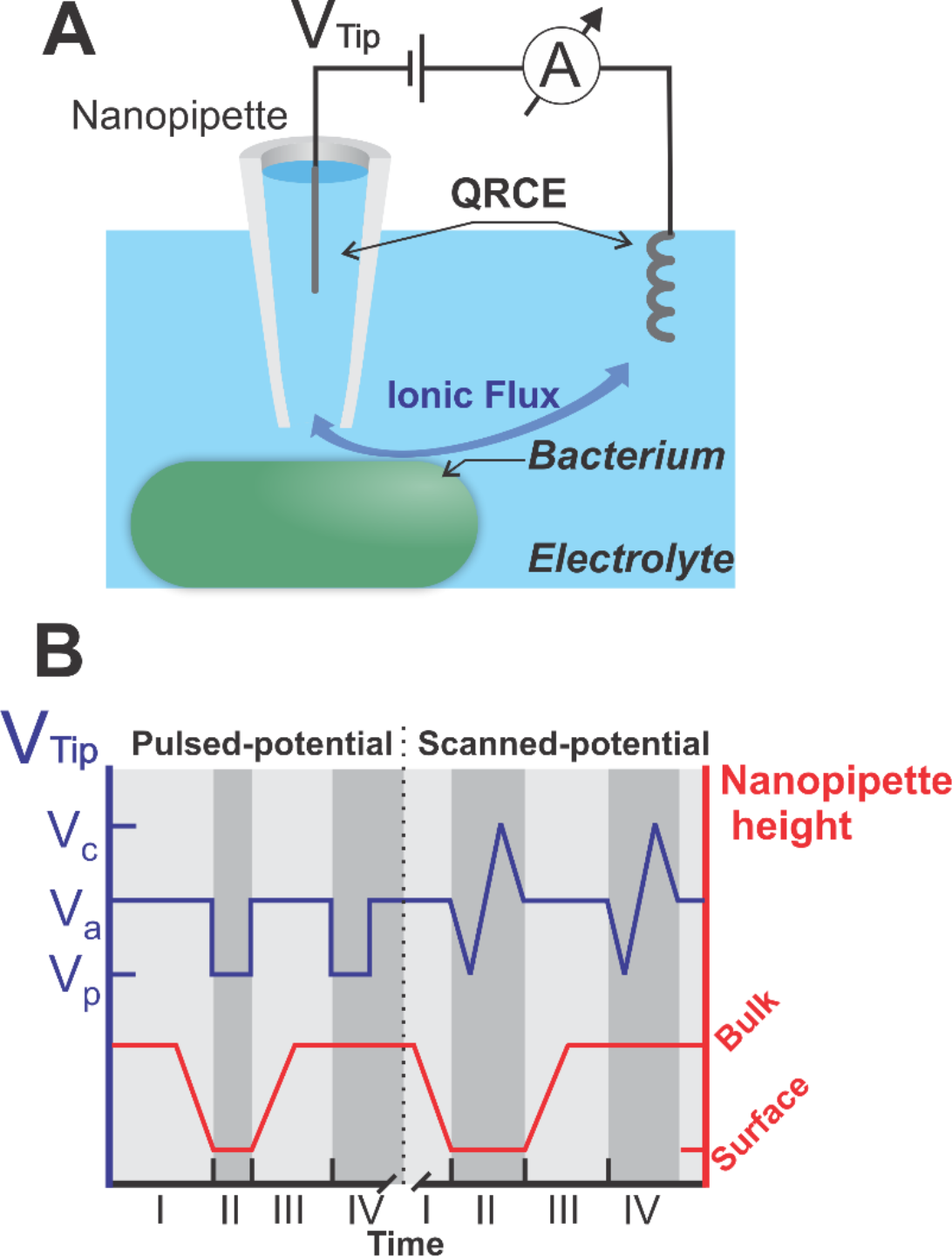
**(A)** SICM schematic depicting a nanopipette probe in electrolyte and connected across two QRCEs, where V_Tip_ is the potential applied to the nanopipette QRCE with respect to the QRCE in bulk solution. **(B)** Infographic to show nanopipette height in red (right y-axis), alongside V_Tip_ (in blue, left y-axis) as a function of time, as applied at each position in an SICM map. Two different V_Tip_-time profiles were applied, as shown on the left and right halves of the graph; pulsed (left) and scanned-potential (right). The V_Tip_ potentials defined are: V_a_, the approach potential; V_p_, the pulsed-potential (and one limit of the triangular sweep); and V_c_, the other limit of the linear cyclic potential sweep. For most experiments, V_a_ = 50 mV, V_p_ = −500 mV, and V_c_ =500 mV. See more information in the text.

### FEM Simulations

2D axisymmetric models of the nanopipette in bulk solution and near the substrate were constructed in COMSOL Multiphysics (v. 5.4) with the Transport of Diluted Species, Laminar Flow and Electrostatics modules. For converting experimental nanopipette currents to surface charge values, the real nanopipette geometry, acquired by EM, and experimental conditions, were used in the FEM models. All other simulations exploring the influence of the substrate properties on the nanopipette current response were performed utilizing a representative (average) nanopipette geometry with a lumen radius of 95 nm. Details and models are provided in SI-6.

## Results and Discussion

### Probing the gram-negative cell envelope with pulsed-potential SICM

Pulsed-potential SICM was applied to map live *E. coli* cells adhered to PLL (SI-3) in 50 mM KCl electrolyte. Typical results are shown in Figure 3, where (A) shows the topographical map of the bacterium, which is approximately 2 μm in length and 1 μm in height, in agreement with the literature values and the scanning electron microscopy images taken of cells at this growth stage (SI-7).^40^ In the pulsed potential stage, normalized nanopipette current values were in the range of 0.97-1.05 (Figure 3B). The measured current values were converted to surface charge values using a FEM model (SI-6) that treated the bacterial wall surface as a planar charged impermeable insulator, reflecting the exterior membrane of gram-negative bacteria (Figure 1).^19^ This modelbased conversion resulted in a negative surface charge density of −80 to −150 mC m^−2^ over the surface of the *E. coli* cells (Figure 3C), and a positive surface charge of +30 to +50 mC m^−2^ over the PLL. Interestingly, a significant degree of variation of the surface charge can be observed over the bacterium, where the central apex of the bacterium generates a higher normalized current (and therefore a more negative charge) than at its edges (Figures 3B and C).

**Figure 3:**
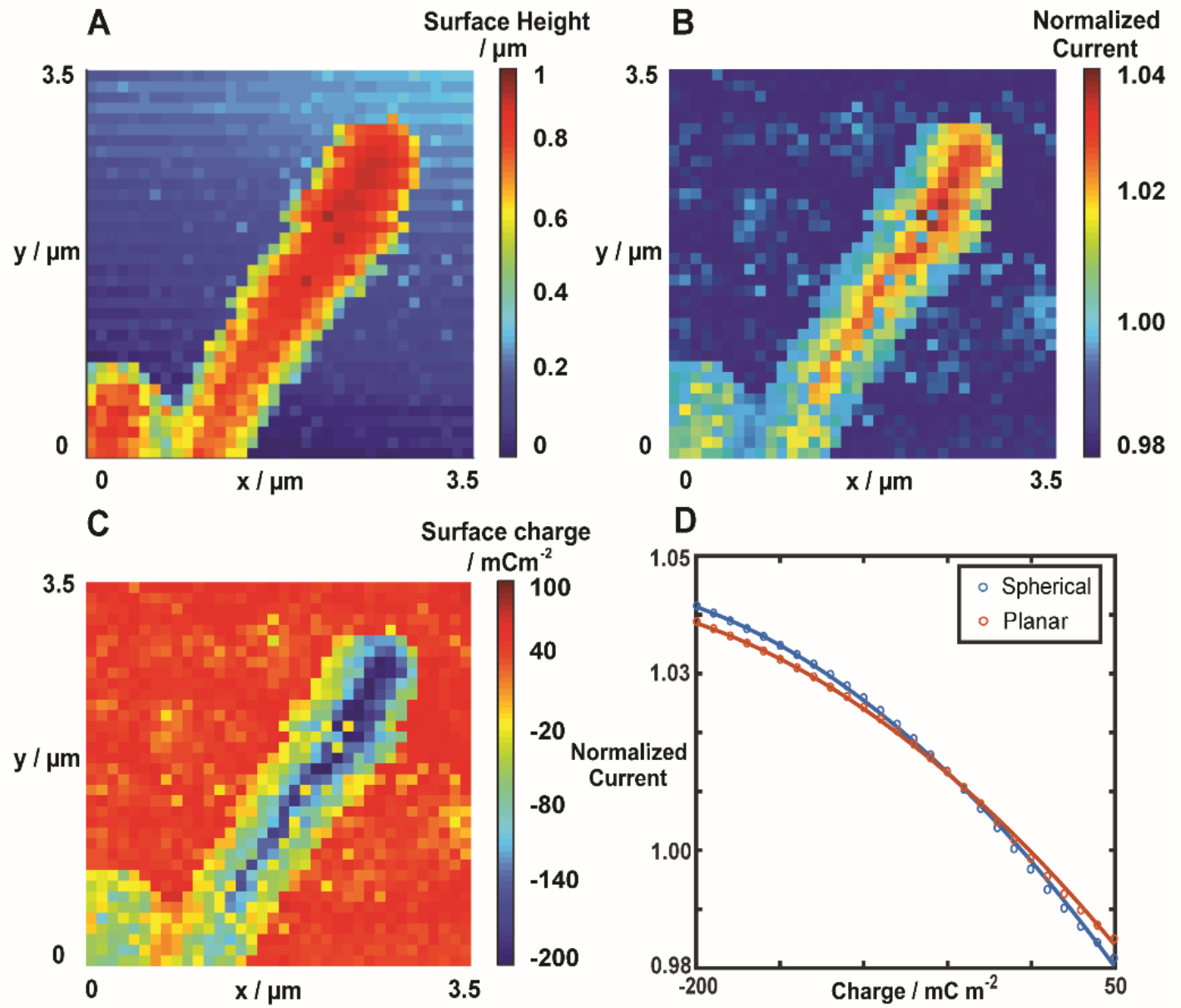
SICM topography **(A),** normalized current **(B),** and local charge density (mC m^−2^) following FEM simulation **(C)** maps of E. coli collected concurrently. **(D)** Comparison of the FEM model simulation results using a planar (red) and spherical (blue) cell form (see SI-6). Lines are fitted calibrations through the simulated points. SICM experiment conditions: 50 mM tris buffered KCl at pH 7, bacteria adhered to PLL. V_p_ = −500 mV and V_a_ = 50 utilizing a 2 % feedback current threshold (see Methods). Nanopipette inner radius of 93 nm, with 100 nm hopping steps (pixel separation). Images are raw data with no interpolation.

While the observed differences in current between the apex and edges of the bacteria are clear, as seen in Figure 3B, the corresponding surface charge differences could be affected by the model-based conversion of the experimental data. To explore this possibility, we modeled the charged surface as both a sphere (radius 0.5 μm, commensurate with that of the bacterium) and a planar surface, with the nanopipette set axisymmetrically above in both cases (see SI-6 for more details of the models). In the corresponding FEM simulations using these two different models, the nanopipette-substrate distance was set for the same current set point (2 % decrease in current from the bulk value), *i.e*. the same overall gap resistance. A comparison of the normalized current *vs*. surface charge resulting from the two models is shown in Figure 3D. Overall, the difference between the models is small, with a maximum of *ca*. 0.002 difference in normalized current values at the highest surface charge considered (−200 mC m^−2^). When calibrated to the experimental scan data over the bacterium in Figure 3, the charge value calculated from the planar model was found to be at most *ca*. 10 mC m^−2^ more negative compared to the spherical model (see SI-8), which is negligible compared to the range of values. As the difference between the models is relatively small, and the experimental scans include the flat background substrate, the planar model is used herein when describing the gram-negative bacterial scans. Note that although COMSOL Multiphysics can, in principle, be used for 3-dimensional simulations,^21,41,42^ the accuracy can be compromised and/or simulations become computationally expensive for complex problems, such as here, where there is an interplay of ion migration and electroosmotic fluid flow, and the need to consider charged interfaces. We thus retained a 2D model to assess the effect of a curved interface. The choice of a spherical object to assess the curvature of the substrate also maximizes the effect compared to the real situation where, for example, the majority of the bacterium is approximately cylindrical (Figure 3A). Further evidence that the significant differences in charge between the top and sides of the bacterium are not topographical in origin also comes from the fact that the generally much higher negative charge density (≤ −100 mC m^−2^) at the tops of the bacterium extends right to the very end of the bacterium where there is significant curvature (Figure 3A-C).43

Figure 3 also shows that there are small patches of low negative charge (with values of *ca*. −20 mC m^−2^) within areas of much larger negative charge density (≤ −100 mC m^−2^). We considered whether some of these significant local variations in surface charge could be due to the SICM tip causing potential changes near the bacterial cell well and activating voltage-gated or mechanosensitive ion channels (MSCs). To explore this possibility, we used the experimentally determined nanopipette geometry with FEM simulations to analyze potential field around the nanopipette tip (Figure 4). We find that the majority of the tip-induced voltage drop is in the first few microns within the nanopipette (Figure 4A),^16^ presumably due to the high resistance at the nanopipette lumen (140 MΩ, calculated from *i-V* curves recorded in bulk solution). Although some of the potential field extends from the end of the nanopipette, the impact of this on the bacterial surface, at the highest pulsed nanopipette potential of −500 mV, is expected to be only −15 to −25 mV (Figure 4B), considerably less than the −50 mV depolarization potentials reported for example for *E. coli* K^+^ ion channels.^44^ Unlike voltagegated ion channels, MSCs are sensitive to both potential and mechanical stresses which can arise from external pressure.^45^ We thus further analyzed the FEM simulations to account for the pressure exerted by electroosmotic fluid flow from the nanopipette to the cell surface (discussed in SI-8). We found this to be *ca*. −45 Pa, suggesting a less than 10 % probability of opening MSCs at the interfacial potential applied here.^45^ Together, these additional analyses suggest that most of the observed charge heterogeneity is unlikely due to bacterial channel gating, and more likely to be a genuine heterogeneous charge distribution along the cell wall or a signal from natively functional ion channels.

**Figure 4.**
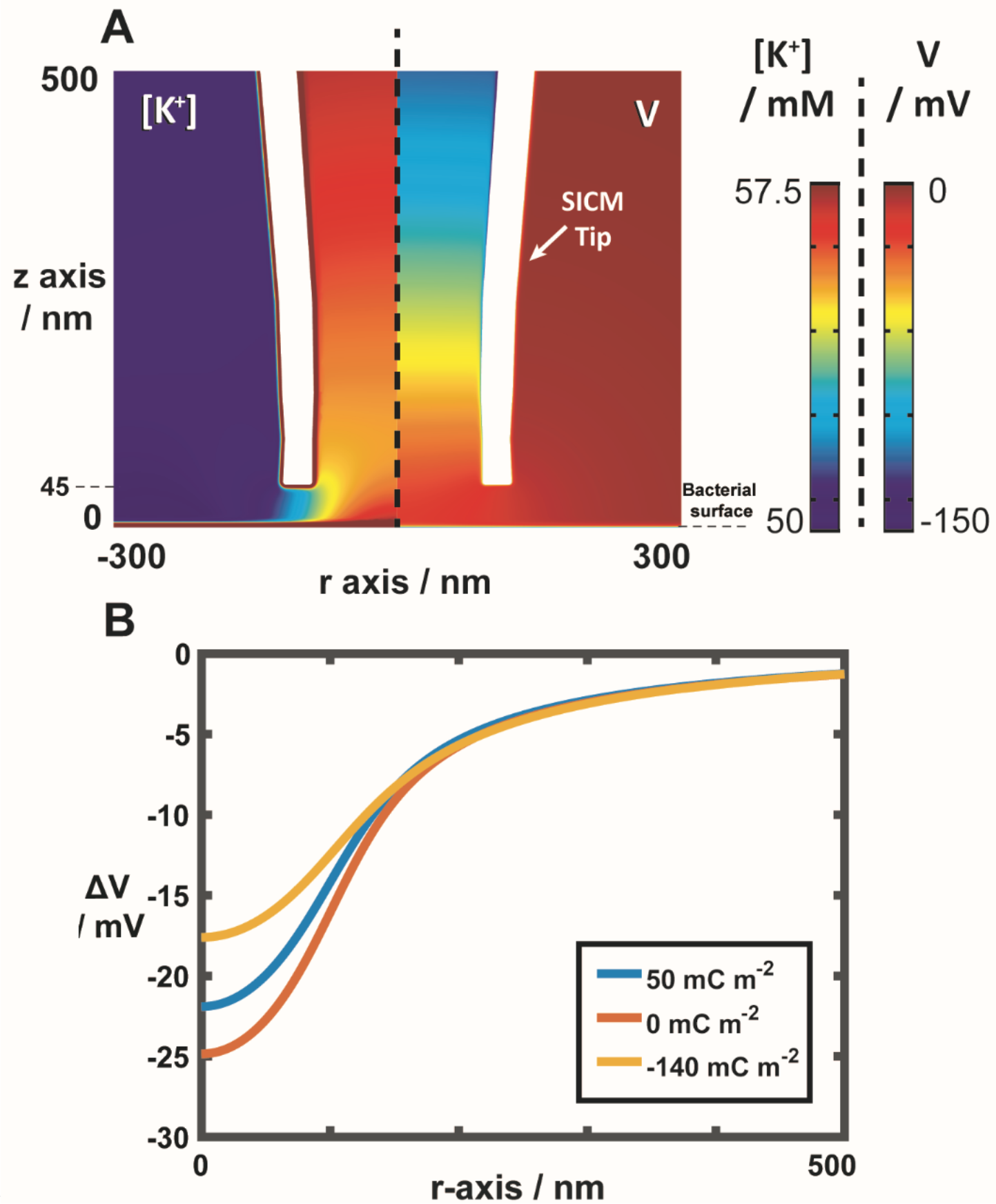
Error! No text of specified style in document.: FEM-based simulation results of pulsed-potential SICM (V_Tip_ = −500 mV after 20 ms) at a charged surface (−140 mC m^−2^). **(A)** Potential and [K^+^] distribution at surface-nanopipette interface: left panel shows the [K^+^] distribution and right panel shows the corresponding potential distribution, vs. bulk QRCE potential. The inner nanopipette radius is 95 nm, and the nanopipette-substrate separation is 45 nm. **(B)** Solution potential with respect to bulk (DV), from the nanopipette center radially outwards at the surface-solution interface (3 surface charge densities as defined), at a nanopipette-surface separation of 45 nm. Values for different surface charge densities are shown (see legend).

### Probing the gram-negative cell envelope with scanned-potential SICM

Scanned-potential experiments were implemented to further confirm the findings of the pulsed-potential SICM measurements across a wider range of potentials, and to further investigate the possible contribution of ion flux from voltage stimulation of cell surface efflux pumps and channels.^46^ In these experiments, the potential at the nanopipette was scanned in a triangular waveform across a range of values and the resulting current recorded, generating *i-V* curves at every position of the physical scan (Figure 2B). While it increases the overall scanning time (*ca*. 15 seconds/position), the use of a scanned-potential protocol provides a greater depth of information (potential-resolved images presented as Movie 1 in the SI, *vide infra*)^19^.recording of the topographica lTo achieve better stability of bacteria over these longer scanning periods, these experiments were done using Cell-Tak, which provides a stronger hold on the bacteria.

Figure 5 provides a summary of typical results from the scanned-potential measurements of *E. coli*. Figure 5A shows *i-V* curves at selected positions over two bacteria; in all cases there is slight current rectification, with larger current magnitude at the extreme negative potential compared to positive potential (+/- 500 mV), demonstrating that the gramnegative bacteria surface responds to potential-scan SICM in a way similar to previous descriptions of surface-induced rectification (SIR).^13,17,19^ The insert in Figure 5A shows negligible current rectification between −50 and 50 mV, demonstrating that the ionic current at the nanopipette is not affected by the surface charge at the approach potential used (50 mV) which allows for consistent nanopipette approach distances throughout the image and the recording of the topographical map in Figure 5B.^15^

**Figure 5:**
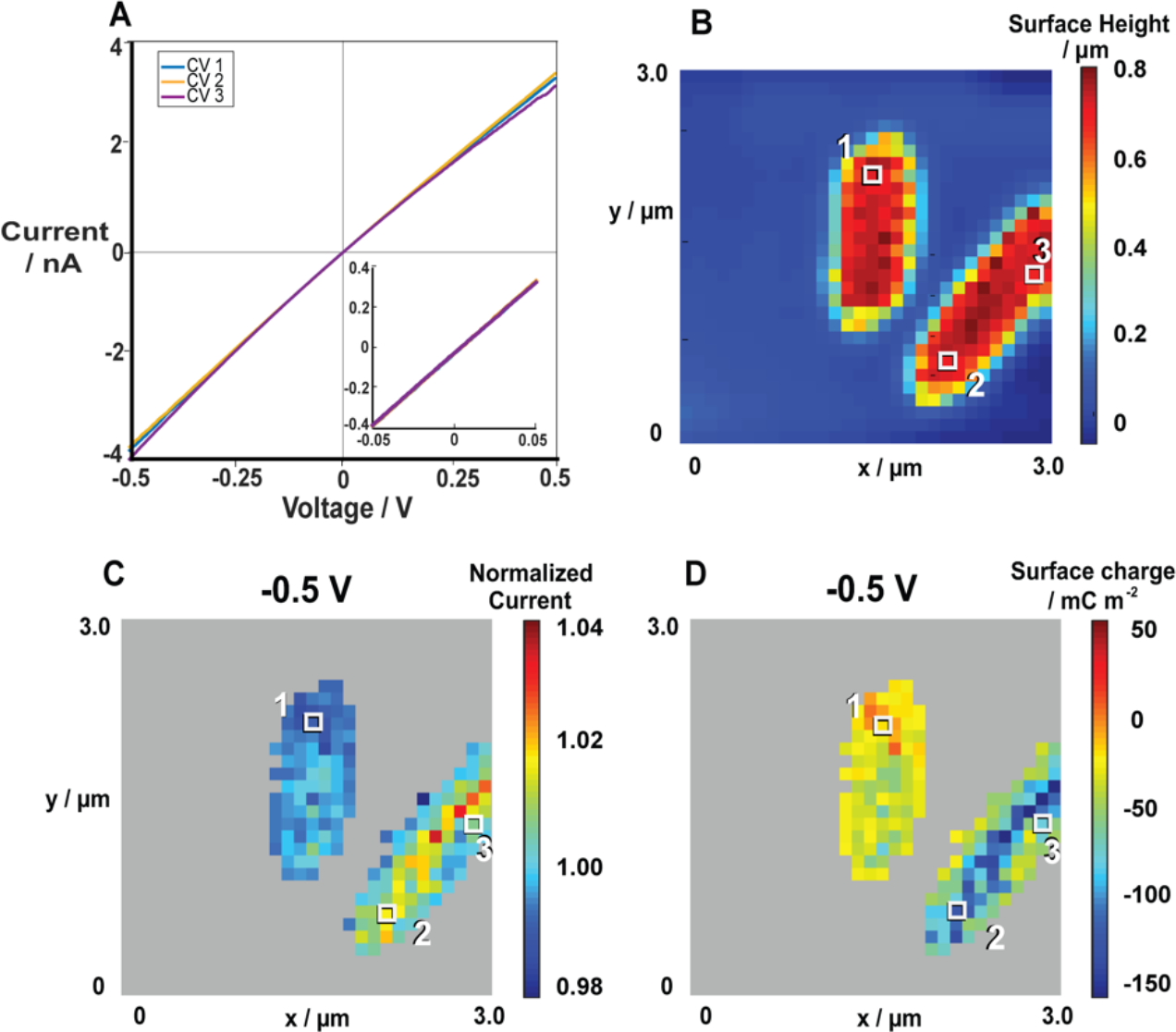
Scanned-potential SICM as applied to E. coli adhered with Cell-Tak in 50 mM tris buffered KCl solution at pH 7. **(A)** i-V curves at selected labelled positions (1-3), with the insert showing a close up of currents between −50 and 50 mV. **(B)** Topography maps of E. coli. **(C)** Normalized current maps when V_Tip_ = −500 mV during the potential scan. **(D)** Corresponding FEM-based charge density values (mC m^−2^) at −500 mV. Nanopipette inner radius of 90 nm, with 100 nm hopping steps (pixel separation).

The normalized current and charge values at the negative extreme (−500 mV) are shown in Figure 5C and 5D, respectively. Normalized current and corresponding surface charge images at the positive potential extreme (500 mV) are given in SI-9, and a movie of the scan showing the normalized currents throughout the potential range (separated, potential resolved images) is presented as Movie 1. The data in Movie 1 confirm the validity of the approach adopted: there is little contrast in the normalized current value between the bacteria and support when the applied potential magnitude is small (insensitive to surface charge) and the contrast develops at extreme potentials (sensitive to surface charge). This is particularly striking as the *E. coli* bacteria were adhered using Cell-Tak (compared to PLL for Figure 3), which was found to have high positive charge values. As a consequence, to facilitate the visualization of the charge distribution over the bacteria cells, normalized currents and charge associated to the substrate (considered to be any normalized current below 0.98, or charge above +50 mC m^−2^) are removed from Figure 5C and 5D. Raw data demonstrating the full range of normalized current and charge is shown in SI-9, where further discussion on Cell-Tak charge properties can be found.

Figure 5D shows the *E. coli* bacteria have a surface charge range between −70 to −120 mC m^−2^, in the same range of values as the pulsed SICM measurements of *E. coli* bacteria shown in Figure 3C. Again, the largest charge density tends to be across the central apex of the cell, and the overall charge density is visually different on the two bacteria, suggesting possible heterogeneities at the individual cell level across a cell population, which requires further analyses in future studies. Overall, there is agreement between the results of the two SICM regimes (pulsed and scanned potential).

As with the pulsed-potential, the scanned potential measurements also indicate local current (charge) heterogeneities on the bacteria surface (Figure 5C/D). Thus, we again briefly consider whether the activation of a single MSC by the SICM measurement could occur at the largest applied nanopipette potentials (Figure 4B). MSCs, when activated, can generate currents between 40-75 pA,^47,48^ well above the noise level of SICM measurements (*ca*. 4pA), and multiple channels could be located within a scanning area, which would lead to higher currents. Thus, the observed high current patches could be natively active MSCs or ion channels. The possibility of SICM-induced activation of channels, however, is not evident; we see no changes in the measured *i-V* profiles (Figure 5A) nor in the potential resolved images (Movie 1), indicating that there is no SICM potential induced activation of channels at larger applied potentials. An interesting application of SICM in the future would be to explore whether conditions could be generated to deliberately activate ion channel function, without physical patch (contact) on the nanopipette.

### Probing the gram-positive cell envelope with pulsed-potential SICM

We now consider experiments performed on the gram-positive *B. subtilis* (using agarose for adhesion). Typical pulsed-potential SICM results are displayed in Figure 6, with SICM topographical mapping (Figure 6A) giving cell dimensions corresponding well with electron microscopy images (SI-7). The normalized currents over the bacteria from the potential pulse (Figure 6B) are significantly higher at 1.08-1.12 compared to the *E. coli* case (0.98-1.05, Figure 3B), for the same experimental conditions, suggesting a high density of stationary negative charge at the interface. Inter-cellular heterogeneity in both cell size and normalized current can be observed in Figure 6, highlighting again the value of single cell measurements.

**Figure 6:**
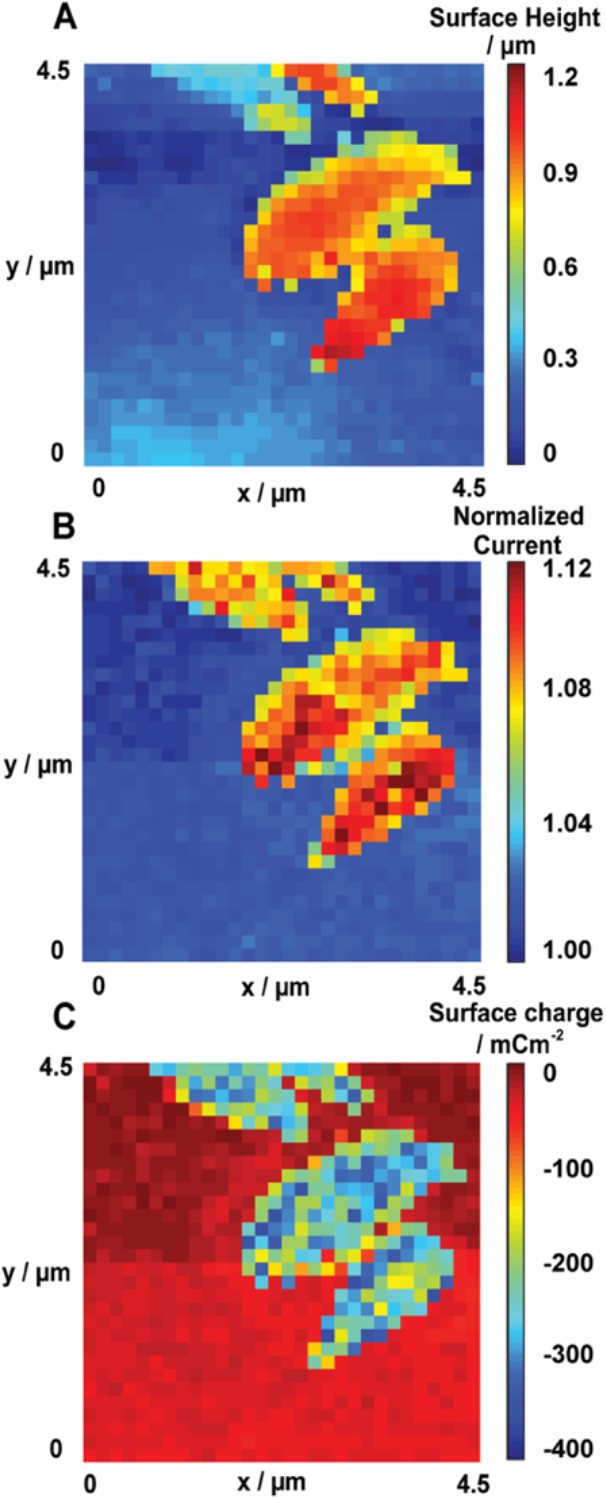
SICM topography **(A)** and normalized current **(B)** of Bacillus Subtilis (Δhag) on an agarose substrate, collected concurrently, along with the resulting charge density values (mC m ^−2^) from the FEM model **(C)**. SICM experiment conditions: 50 mM tris buffered KCl at pH 7. V_p_ = −500 mV and V_a_ = 50 mV utilizing a 2 % feedback current threshold (see Methods). Nanopipette inner radius of 98 nm, with 150 nm hopping steps. Note that panels show raw data without any interpolation.

Using the model outlined in SI (SI-6), surface charge density values were calculated as being between −250 to −350 mC m^−2^, considerably greater than for *E. coli* (*vide supra*), and similar to values in literature from other techniques (*e.g*. electrophoretic mobility) for gram-positive bacteria, where, in low concentration electrolytes, values ranging between −200 and −500 mC m^−2^ have been reported.^49,50^ It is important to note that while the electrolyte composition of 50 mM KCl was tolerated by our bacteria strains (see SI-4), it is possible that the absence of carbon source and physiological media conditions could cause a certain degree of cellular stress, possibly affecting the charge in the bacterial cell envelope.^51^ However, similarly high normalized currents were also found when we used a physiological media (M9m) as the electrolyte to scan *B. subtilis Δhag* (SI-10, Figure S-12).

### Probing the gram-negative cell envelope with scanned-potential SICM

Figure 7 shows a summary of results from scanned-potential mapping of *B. subtilis* cells in 50 mM KCl solution; Movie 2 (SI) shows the full potential scan. The *i-V* curves (Figure 7A) obtained from points above the bacterium or the substrate demonstrate consistent smooth rectification profiles, attributable to SIR,^19,14^ but the similar behavior in a narrow potential range around 0 V, again confirms the validity of the charge mapping strategy: in this potential region the SICM response is insensitive to surface charge and provides a faithful map of topography.

**Figure 7:**
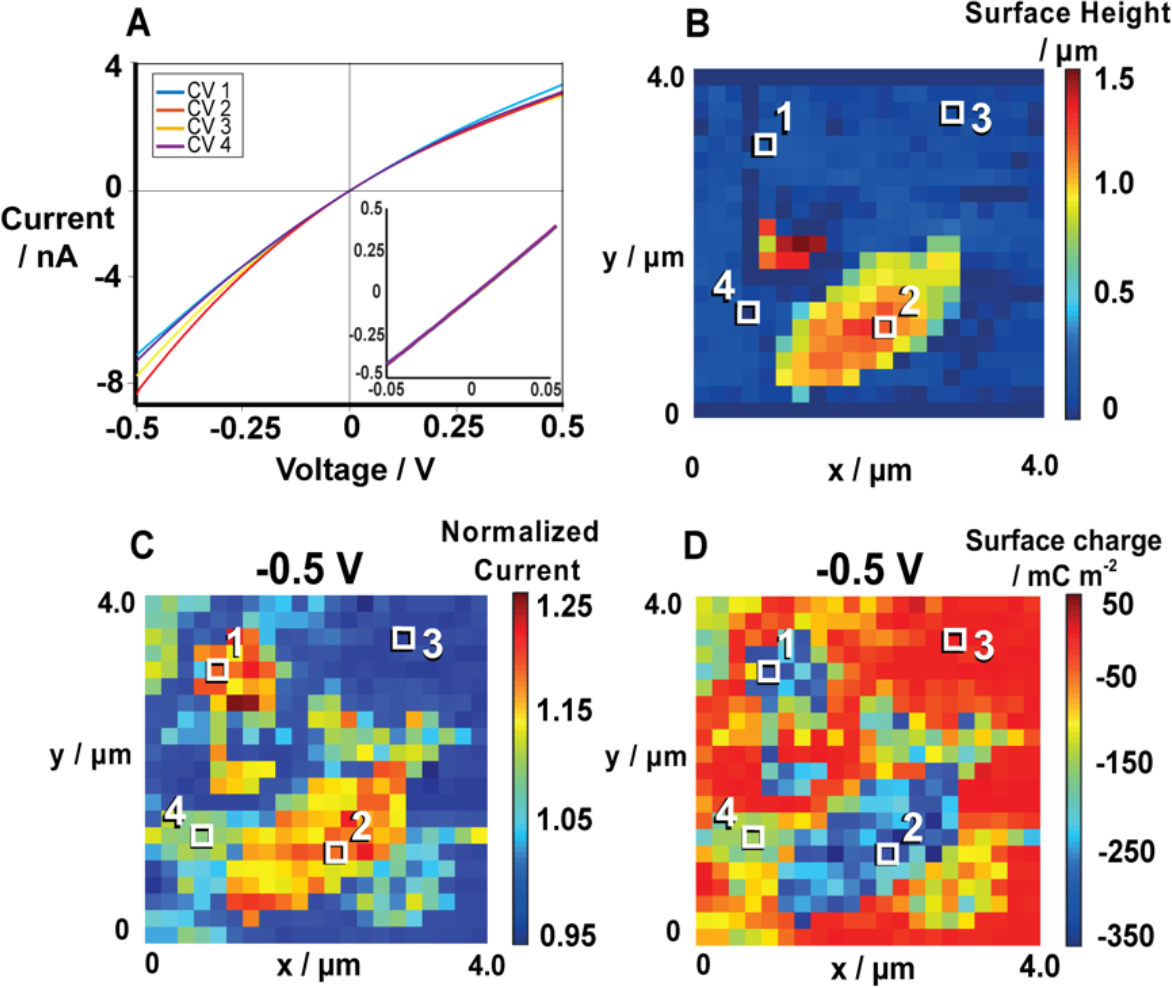
Scanned-potential SICM of B. subtilis (Δhag) in 50 mM tris buffered KCl solution at pH 7, adhered to agarose. **(A)** i-V curves at selected labelled pixels (1-4), with the insert zoomed to between −50 and 50 mV. **(B)** Topography maps of B. subtilis cells. **(C)** Normalized current maps for V_Tip_ = −500 mV. **(D)** Corresponding FEM-based surface charge density values (mC m^−2^) at −500 mV. Nanopipette inner radius of 93 nm, with 200 nm hopping steps. Note that panels B, C and D show raw data without any interpolation.

The topography map (Figure 7B) shows the bacteria distinctly. At *V_Tip_* = −500 mV, the large normalized current ratio (Figure 7C) and the corresponding negative charge density (Figure 7D) over the bacteria is seen. Significantly, however, there is a further region of negative charge density around the bacteria (Figure 7C, D, region labeled 4). We attribute these negative charged regions to the complex ion-permeable matrix known to be secreted by grampositive bacteria, broadly termed the extracellular polymeric substance (EPS).^52,53^ We confirmed this through EM imaging of the bacteria, where a coating was observed over and around the cells (SI-7). EPS consists of polysaccharides, nucleic acids, lipids and polypeptides^54^ and can extend up to a micron from the cell surface,^55^ as found here. Our results here indicate that this material, which is also shown to aid bacterial adhesion,^53^ formation of biofilms,^56^ and nutrient trapping,^57^ is highly negatively charged, and that SICM charge mapping is a powerful means of assessing the regions over which EPS extends.

The difference in the ion current response between the gram-negative *E. coli* and the gram-positive *B. subtilis* could possibly be attributed to differences in the structure of the cell (Figure 1). For *B. subtilis*, the permeable and ion dense peptidoglycan layer is not shielded by an insulating outer membrane and, as such, presents a more accessible ion rich region for the SICM tip response.

### An extended FEM model of the gram-positive cell envelope

Although the charge values recovered from the experimental data using the simplified FEM model are in good agreement with previous reports for both bacterial strains, the apparently very high charge densities for *B. subtilis* require an extended FEM model, based on a more realistic physical description of the cell wall and, in particular, the ion permeable and ion dense peptidoglycan layer. Here, we developed such an extended model specifically to account for the specific aspects of the gramnegative cell envelope. The primary biophysical factors implemented in this extended model, treating the cell as a sphere are: the fixed charge concentration in the peptidoglycan layer (*ρ_f_/F*), where *ρ_f_* is the negative charge concentration and *F* is Faraday’s constant, the thickness of the peptidoglycan cell wall (*t_Wall_*, see SI-6.2) and the mobility of counter ions within it (*μ_wall_*), relative to that in solution. It was previously found that co-ions are excluded from the peptidoglycan cell wall of *Bacillus brevis* at electrolyte concentrations similar to those employed in our experiments,^31^ and hence the model assumed no Cl^−^ partitioning to the cell wall. A full description of the extended model can be found in Sections SI-6 and SI-11.

Using this extended model we re-simulated the SICM experiments as applied to *B. subtilis*, with results shown in Figure 8. Figure 8A shows the concentration profile of K^+^ ions in the vicinity of the nanopipette end and cell wall. At the approach potential (*Va* = 50 mV), [K^+^] is more or less uniform, except in the double layer of the charged interfaces. With a negative pulse potential (*V_p_* = −500 mV) K^+^ is drawn from the cell wall into the nanopipette, leading to an increase in [K^+^] in the vicinity of the nanopipette end. The corresponding potential distributions for these cases are shown in Figure 8B. This is shown in more detail in Figure 8C, where the potential distribution over the thickness of the cell wall and just into the solution (*z*-coordinate) is shown for the cases of no applied potential, and *V_Tip_* set at 50 or – 500 mV. In all these cases, the interior of the cell wall (−34 to 0 nm) reaches the Donnan equilibrium, noted by the plateau in the potential value. The potential difference between the cell wall and nanopipette end (*ΔE* in Figure 8C) are −17.5 mV for the unperturbed case (*V_Tip_* = 0 mV), −19 mV when the approach potential is applied (*V_Tip_* = 50 mV), and −9.4 mV under the pulsed potential (*V_Tip_* = −500 mV). A summary plot indicating how the normalized nanopipette current varies with wall charge density (*ρ_f_ /F*) and ionic mobility within the cell wall (*μ_Wall_*), with all other parameters fixed to literature estimated values (SI-6) is shown in Figure 8D.

**Figure 8:**
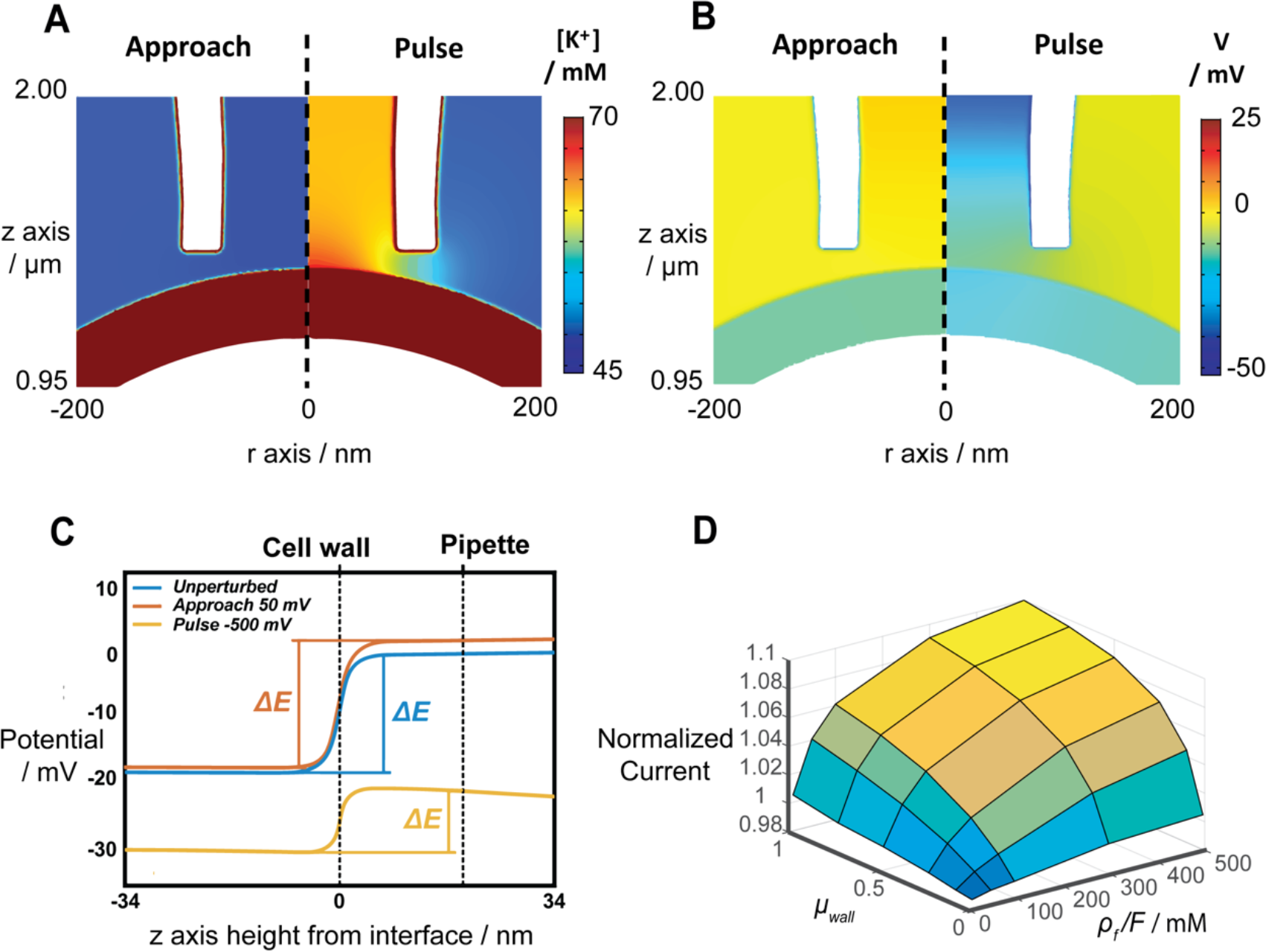
FEM simulations of pulsed-potential SICM at the gram-positive cell surface with an assumed stationary charge concentration (**ρ_f_ /F**) of 100 mM in the peptidoglycan layer. The other parameters were: peptidoglycan cell wall (substrate) thickness of 70 nm; nanopipette-substrate distance of 20 nm; dielectric relative permittivity of the cell wall of 20; and relative ionic mobility within the cell wall compared to solution (μ_Wall_) of 1. **(A)** [K^+^] and **(B)** potential distribution, at the approach potential (50 mV, left) and the pulse potential (−500 mV, right). **(C)** Potential-distance profiles across the cell wall-solution interface (at the cylindrical symmetry axis): with no perturbation (blue); at the approach potential (orange, V_Tip_ = 50 mV); and at the pulse potential (yellow, V_Tip_ = −500 mV). For these plots, the cell wall-interface is at 0 nm, with negative values of z into the cell wall and positive values into the solution (and positions of the cell wall and nanopipette marked by the vertical dashed lines). **(D)** Normalized SICM current (at nanopipette potential of −500 mV) as a function of μ_Wa11_ and ρ_f_/F.

These simulation results show that the normalized SICM current values are dependent on *ρ_f_ /F* and *μ_wall_*, as expected. With increasing charge density and/or the mobility of ions in the peptidoglycan cell wall, the normalized current ratio increases, reaching a maximum value of *ca*. 1.1, close to that observed experimentally for the *B. subtilis* cells over most of the cell surface (Figure 6 and 7). Modifying the cell wall thickness within the range observed by EM (Figure S-13B) causes only minor changes in the simulated normalized currents, because the perturbation caused by the SICM probe only extends into the outermost region of the cell wall (Figure 8B), as also seen with permeable abiotic substrates.^34^

An interesting avenue for future exploration would be to analyze in more detail the current-time transients in pulsed-potential measurements, and apply double pulses, to transiently drive the cell wall (substrate)-interface out of equilibrium with SICM, and monitor its return to equilibrium. Analysis of similar scanning electrochemical microscopy studies has enabled the relative diffusion coefficient and partition coefficient to be determined independently in a single measurement, without the tip contacting the sample.^58,59^ To ensure sensitivity to the cell wall, it may also be beneficial to explore the use of differential concentration SICM, with a different concentration of electrolyte in the nanopipette to that in the bulk solution.^38^

## Conclusions

Here, we demonstrated the application of SICM to the study of the ionic environment of live gram-positive and gram-negative bacteria. Several methods are used for adhering bacteria to the glass substrate allow SICM whilst maintaining cell viability. The adherents selected were not shown to affect the bacteria ionic environment, and similar SICM current responses were seen for the same strains across different substrates. Using FEM simulations to model the influence of the SICM nanopipette tip on the cell wall interface, we were able to understand and account for the potential and hydrodynamic pressure perturbations applied to the bacteria, showing that the SICM tip, at the conditions employed here, can be considered non-inductive to cell physiology, allowing surface charge of both strains to be analyzed using SICM and FEM.

In-line with previous bulk measurements, we found that both gram-positive and - negative bacteria display negative surface charges, with gram-positive *B. subtilis* having a significantly more negative charge than the gram-negative *E. coli*. This was confirmed by using two SICM regimes, a pulsed-potential and a scanned-potential regime, both of which yield similar charge results. The scanned-potential regime also produced similar charge values at both potential extremes, demonstrating that the direction of polarity did not considerably influence the calculated surface charge. The application of FEM simulations allowed us to calculate surface charge for both bacterial types, and an extended model allowed first insights into capturing impact of peptiglycan layer on SICM measurements done on gram-positive bacteria. This extended model has potential to be developed further to capture possibly complex transport phenomena at the gram-positive cell wall.

By utilizing small nanopipettes as the SICM tip we were able to visualize charge heterogeneity between and across the individual bacteria and the charge distribution over the secreted EPS layer. This high spatial resolution, coupled with an extended model of the gramnegative bacteria cell wall allowed initial insights to the intricate interplay between the biophysical and SICM parameters. We also show that SICM has the potential to be applied in a more intrusive manner to investigate ion channels within the cell wall without destroying the integrity of the cell structure, a technique that we envisage could be similar to a form of noncontact patch clamping. Thus, the presented methodology will pave the way to a more thorough understanding of the interconnection between cellular physiology and bioelectrical microenvironment of cells, benefiting a broad range of research areas including cell biology, bacterial adhesion, antibiotic resistance, and biofilm formation, and making a significant impact on life science research and development.

## Author Contributions

Methods were designed by KC, BJ and GNM, with experiments performed by KC and BJ. Simulations were developed by JT, GNM and DP, and performed by KC and JT. CZ and MA provided support and advice for biological materials, protocols and analysis. PRU and OSS conceived the study, supervised and helped with data interpretations. The manuscript was written by KC, GNM, JT, OSS and PRU.

## Acknowledgements

JT and KC thank the EPSRC for support through MAS CDT, grant number EP/L015307/1. BJ thanks EPSRC/Unilever for an iCASE award. GNM acknowledges support from the European Union’s Horizon 2020 research and innovation programme under the Marie Skłodowska-Curie grant agreement 790615 (FUNNANO). PRU thanks the Royal Society for support through a Wolfson Research Merit Award. The authors would all like to acknowledge the support of the Bio-Electrical Engineering Innovation Hub, University of Warwick, funded by the UK’s Biological and Biotechnological Sciences (grant no. BB/S506783/1) and Engineering and Physical Sciences Research Councils. The authors acknowledge the Kearns lab, from Indiana University, who developed the *B. subtilis* mutants used that we obtained through Munehiro Asally. We also gratefully acknowledge Corinne Bailey and Saskia Bakker, from the University of Warwick Advanced Bioimaging platform, for assistance with the EM imaging of the bacterial strains, and Ian McPherson for fruitful discussions. Finally, we wish to acknowledge Teuta Pillizota, from the University of Edinburgh for fruitful discussions and support regarding adhesion of bacteria to substrates.

