## Supplementary Information for "Scanning ion conductance microscopy reveals differences in the ionic environments of gram positive and negative bacteria"

### Contents

**SI-1 Bacterial strains .......... S.3**

**SI-2 Bacteria culturing .......... S.3**

**SI-3 Sample preparation .......... S.6**

**SI-4 Determining viability of bacterial strains using optical density ($OD_{600}$) .......... S.8**

**SI-5 Nanopipette fabrication and characterization .......... S.10**

SI-5.1 Nanopipette fabrication .......... S.10

SI-5.2 Nanopipette characterization .......... S.10

**SI-6 Finite element method (FEM) simulations .......... S.11**

SI-6.1 Simple FEM model for bacterial charge .......... S.12

SI-6.2 Extended FEM model for gram-positive bacteria .......... S.14

SI-6.3 FEM Simulation framework .......... S.18

**SI-7 Electron microscopy analysis of bacteria .......... S.18**

**SI-8 Further analysis of pulse-potential *E. coli* scan .......... S.21**

**SI-9 FEM simulations for the scanned-potential *E. coli* scan .......... S.24**

**S-10 SICM of *B. subtilis* in biological media .......... S.26**

**SI-11 Further results from the extended FEM model .......... S.27**

#### SI-1 Bacterial strains

The bacteria used in this study were the wildtype *Escherichia coli* K12 strain and a *Bacillus subtilis* NCIB 3610 mutant strain deplete of the *hag* gene. The former was originally obtained from the German Culture Collection Centre (DSMZ) (DSM No. 498), while the latter strain (denoted as DS1677) is referenced in Mukherjee *et al.* (2013).[1] The Δ*hag* mutant lacks production of flagellin (termed as Hag), which is a key subunit in the bacterial flagella. Therefore, species deplete in Hag have limited flagella, resulting in reduced motility. For the application of SICM, the limited motility enabled sample preparation for the SICM scan (see below for further details).

#### SI-2 Bacteria culturing

Bacteria were grown in a modified M9 media (M9m), based on the established minimal M9 media,[2] but adapted for growing a range of bacteria of interest including the *E. coli* and *B. subtilis* used in this work. The media is also adapted to render it suitable for use as an electrolyte for SICM experiments. All reagents were of analytical grade and used without further purification. Deionized water (Milli-Q, resistivity ca. 18.2 MΩ.cm at 25 °C) was used in the preparation of all solutions. Listed in Table S-1 to S-4 are the final components included in 1 L of the M9m with glucose as a carbon source. All media were adjusted to pH 7, autoclaved, and used within a week of preparation.

**Table S-1:** Full M9m media components.

| | Details explained in | Components added to 1 L |
|---|---|---|
| 5x M9 minimal salts | Table S2 | 200 mL |
| 1 M $MgSO_4 \bullet 7H_2O$ | - | 2 mL |
| Carbon source (0.4 % w/v) | - | 20 mL |
| 1 M $CaCl_2$ | - | 100 µL |
| 10 mM $FeCl_3 \bullet 6H_2O$ | - | 1 mL |
| 1000x Trace metal solution | Table S3 | 1 mL |
| 5-vitamin solution | Table S4 | 1 mL |

**Table S-2:** Quantities and concentrations for M9m minimal salts.

| | **Components added for stock solution 1 L** | **Concentration in 1 L stock (mM)** | **Concentration in 1 L final full M9 media (mM)** |
|---|---|---|---|
| $Na_2HPO_4 \bullet 7H_2O$ | 64 g | 238 mM | 47.6 mM |
| $KH_2PO_4$ | 15 g | 110 mM | 22 mM |
| NaCl | 2.5 g | 43 mM | 8.6 mM |
| $NH_4Cl$ | 5 g | 93 mM | 18.6 mM |

**Table S-3:** Quantities and concentrations for 1000x trace metal solution.

| | **Components added to 1 L stock solution** | **Concentration in 1 L stock (mM)** | **Concentration in 1 L final full M9 media (μM)** |
|---|---|---|---|
| $CuCl_2 \bullet 2H_2O$ | 5.455 mg | 0.032 mM | 0.032 μM |
| $ZnSO_4 \bullet 7H_2O$ | 219.968 mg | 0.765 mM | 0.765 μM |
| $CoCl_2$ | 21.436 mg | 0.169 mM | 0.169 μM |
| $Na_2MoO_4 \bullet 2H_2O$ | 399.218 mg | 1.650 mM | 1.650 μM |
| $H_3BO_3$ | 2862.729 mg | 46.300 mM | 46.300 μM |
| $NiCl_2 \bullet 6H_2O$ | 998.382 mg | 4.200 mM | 4.200 μM |
| $Na_2WO4 \bullet 4H_2O$ | 80.154 mg | 0.243 mM | 0.243 μM |
| $Na_2SeO_3 \bullet 5H_2O$ | 59.966 mg | 0.228 mM | 0.228 μM |

**Table S-4:** Quantities and concentrations for 5-vitamin solution.

| | **Components added to 100 mL stock solution** | **Concentration in 100 mL stock (mM)** | **Concentration in 1 L final full M9 media (μM)** |
|---|---|---|---|
| Biotin | 2 mg | 0.082 mM | 0.082 μM |
| Pyridoxine hydrochloride | 10 mg | 0.486 mM | 0.486 μM |
| Thiamin hydrochloride | 5 mg | 0.148 mM | 0.148 μM |
| Riboflavin | 5 mg | 0.133 mM | 0.133 μM |
| Nicotonic acid | 5 mg | 0.406 mM | 0.406 μM |

#### SI-3 Sample preparation

For consistent SICM measurements, cells had to be stationary during the scanning process. During the optimization stage of the experimental methodology, several adhesives were investigated for their ability to adhere bacteria to the cover glass bottomed sample dishes. Those used in the SICM experiments included in this paper are described below.

**Thin agarose layers**

These agarose layers were poured and spread thinly, thus allowing the bacteria to be visualized atop the agarose using the high magnification lens on an inverted microscope, where the optical capabilities facilitated the positioning and tracking of the nanopipette over areas of interest. Based on the agarose pads method by Young *et al.* (2011),[3] agarose solutions were made using 0.8 % (w/v) Cleaver Scientific low melt agarose (CSL-LMA100) in artificial seawater media (ASWm) basal salts and 50 mM sodium acetate (as described in Zerfaβ *et al.*, 2019).[4] The solution was autoclaved for 30 minutes to melt the agarose whilst maintaining sterility. The agarose solution flask was transferred to a water bath at 50 °C to keep the agarose liquefied.

For sterility, the agarose layers were prepared inside a microbiological safety cabinet. 450 μL of agarose solution was pipetted in an outwards expanding spiral from the center of a 50 mm cover glass bottomed dishes (WillCo Wells, USA, HBST-5040, glass thickness approximately 170 μm), the dish was swiftly rotated to evenly spread the agarose and then placed in the laminar flow hood until set, creating a uniform layer. From the volume of agarose solution added to the dish, the agarose layer thickness was calculated to be approximately 0.25 mm. Solidified agarose plates were sealed and kept at 4 °C in a fridge until use (but never for more than one week).

Prior to SICM experiments, 100 μL of an overnight culture (optical density at 600 nm ~ 0.3) was pipetted dropwise across the agarose layer, and rotated to evenly spread. The sample was then incubated for 30 minutes at room temperature. Following this, the dish was placed into an incubator at 37 °C for 30 minutes to dehydrate the agarose and evaporate any residual liquid. Whilst it is possible to adapt this method to incorporate other media instead of the ASWm, due to the transparency and minimal crystallization, clouding or decolorization, the ASWm pads had the best optical properties and stability.

**Poly-L- lysine**

As evidenced by Wang *et al.*,[5] poly-L-lysine (PLL) has minimal effect on bacterial viability if the substance is adhered to a substrate. In order to adhere bacteria using PLL, 50 mm cover glass bottomed dishes (glass thickness approximately 170 μm) were coated in 500 μL of 0.01 % PLL (Sigma Aldrich, sterile-filtered) for 15 seconds. The PLL solution was then extracted via pipette and the dish washed 5 times with 1 mL applications of de-ionized (DI) water. This procedure leaves behind a thin transparent surface layer of PLL. An overnight bacteria culture (50 μL, $OD_{600}$ approximately 0.8) was added to the dish, which was rotated to evenly coat the surface. The sample was then allowed to incubate for 30 minutes at room temperature. The dish was then washed 3 times with application of 1 mL DI water to remove any un-adhered bacteria.

**Corning Cell-Tak**

Cell-Tak was used in accordance with the adsorption method provided in the instructions by Corning (Catalog 354240, 354241). Cell-Tak was purchased at 2.36 mg $mL^{-1}$ concentration, in acetic acid. For coating each 50 mm diameter cover glass bottomed sample dish, 7 μL of Cell-Tak was added to 193 μL of sterile 100 mM sodium bicarbonate solution, adjusted to pH 8 with 5 % acetic acid, and vortexed for 10 seconds. The solution was then immediately added to the center of the dish and spread to cover approximately two thirds of the dish base. The dish was placed in a 30 °C incubator for 20 minutes, then washed with MilliQ water to remove residual sodium bicarbonate. 50 μL of an overnight bacterial culture

($OD_{600}$ approximately 0.8) was added to the middle of the dish and swirled to cover the Cell-Tak layer. The sample was left for 30 minutes at room temperature allowing adherence of cells prior to SICM.

#### SI-4 Determining viability of bacterial strains using optical density ($OD_{600}$)

All of the SICM experiments were run in an aqueous 50 mM KCl electrolyte. To ascertain that this solution was suitable for the cells, cultures of *B. subtilis Δhag*, and *E. coli* WT were grown overnight in M9m at 37 °C. A 1 mL sample of each culture was centrifuged for 5 minutes at 5000 rpm to sediment the cells, where the supernatant was then removed and replaced with 1 mL of a 50 mM KCl solution buffered to pH 7 with Tris. The cells were resuspended and left in the microbial flow hood at room temperature for 8 hours, a similar maximum length of time as in SICM experiments – note that most SICM experiments took much shorter than this maximum time. A control condition was included, where sedimented cells were instead suspended in fresh M9m, to determine if the centrifugation process affected the cell viability following incubation. After incubating for 8 hours, all cultures were centrifuged for 5 min, at 5000 rpm, and resuspended in 10 mL of fresh M9m (the 10-fold dilution of cells in media was used to ensure that a representative proportion of the incubated cells entered to growth plates, so as to highlight any difference between KCl-incubated and M9m-incubated cells). 8 replicates of 200 μL samples from each condition were added to a transparent 96 well plate (Falcon 96 Well Cell Culture plate, sterile (Corning, UK, #353072)), along with a blank M9m for background subtraction. The optical density at 600 nm ($OD_{600}$) was measured using a BMG Clariostar plate reader, set at 37 °C with a double orbital shaking at 150 rpm. Growth measurements were taken once every 10 minutes for 2000 minutes. The results are shown in Figure S-1.

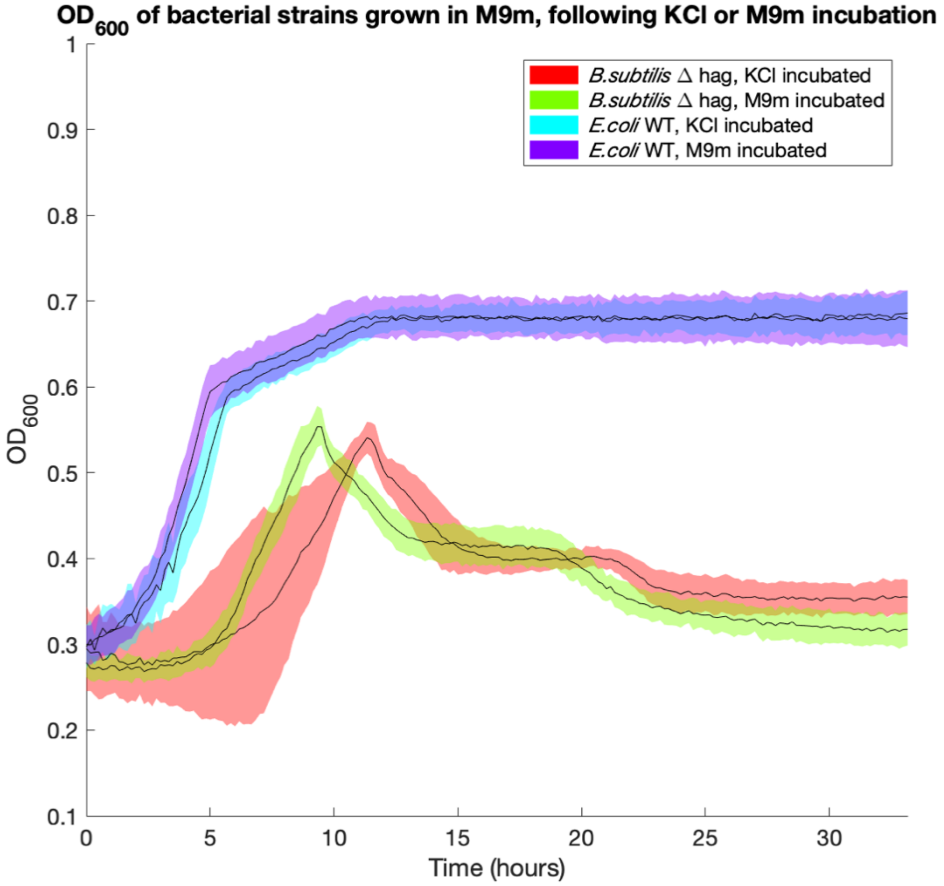


***Figure S-1:*** *Mean (n=8) optical density at 600 nm ($OD_{600}$) of different bacterial strains as grown in M9m media, following 8 hours incubation in either 50 mM KCl or full M9m. Plate reader measurements were made across 33 hours, one measurement every 10 minutes. Lines are used to guide the eye, and standard deviation is shown by the shaded areas.*

Even with the greater standard deviation, the *B. subtilis* strains incubated in KCl show a more gradual exponential growth, compared to M9m incubated cells. This could imply some cell death in the KCl condition that lowers the numbers at the initiation of the plate reader recording, or the exposure to KCl causes a delay or recovery in the exponential growth. However, incubation in M9m could increase cell growth and result in a higher initial viable cell concentration at the initiation of the plate reading. There is also a slight delay in the exponential growth in *E. coli*, but this delay is less prominent.

In conclusion, these results do show a difference in growth between the KCl- and M9m-incubated condition for all strains, however exponential growth still peaks within a reasonable time frame following inoculation. This indicates that a considerable number of cells are still viable following KCl incubation.

#### SI-5 Nanopipette fabrication and characterization

##### SI-5.1 Nanopipette fabrication

Nanopipettes were pulled from borosilicate glass capillaries (o.d. 1.2 mm, i.d. 0.69 mm, Harvard Apparatus) using a laser puller (P-2000, Sutter Instruments: pulling parameters 200 nm pipettes: Line 1: Heat 330, Fil 3, Vel 30, Del 220, Pull -; Line 2: Heat 300, Fil 3, Vel 40, Del 180, Pul120).

##### SI-5.2 Nanopipette characterization

Pulled nanopipettes were characterized regarding their inner radius and overall probe geometry (important for the FEM simulations) by scanning electron microscopy (SEM) using a Zeiss Gemini 500 SEM (operating in scanning transmission electron microscopy (STEM) mode). STEM images of typical nanopipettes used in the SICM experiments are shown in Figure S-2 (lumen diameter approximately 180 nm).

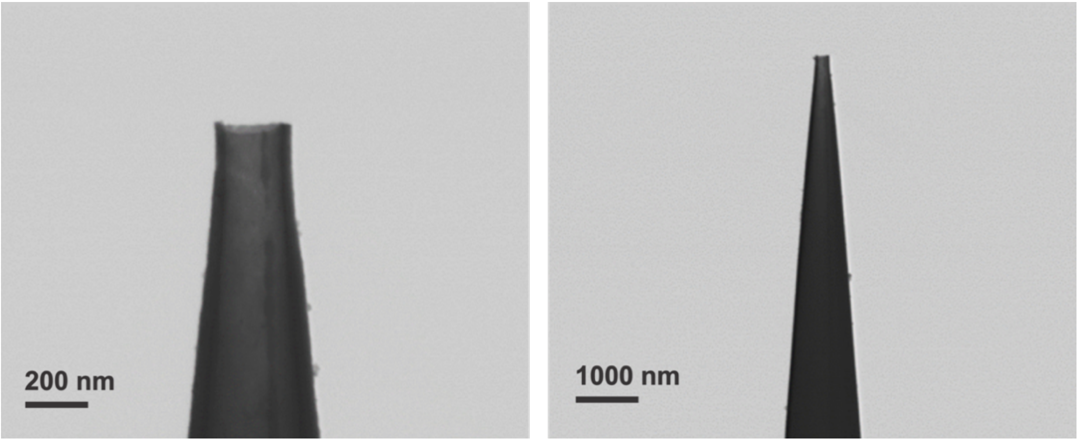


***Figure S-2:*** *Typical STEM images of a nanopipette used in SICM.*

#### SI-6 Finite element method (FEM) simulations

FEM simulations were constructed in COMSOL Multiphysics (v5.4). Bacteria studied here are rod shaped. While a 3D COMSOL model would be necessary to account for the full shape, the additional dimension results in a considerable increase in complexity and computational time (especially for the nanopipette for which mass transport is complex, *vide infra*), hence for the scope of this work a 2D model was used. To account for potential differences in calculated charges due to topography, 2D axisymmetric spherical and planar models (for the substrate surface) were compared.

The electrostatics, transport of diluted species, and laminar flow modules of the COMSOL package were used to model the experimental system. In these models a 2D axisymmetric domain represents the SICM nanopipette, using dimensions taken from STEM images of the nanopipettes (*e.g.* Figure S-2). The nanopipette walls had a surface charge density of -30 mC $m^{-2}$ applied in all simulations, reasonable for borosilicate glass in aqueous solution under the experimental conditions.[6] Experimental *i-V* curves measured in bulk solution were compared against the simulated *i-V* curves resulting from the models to confirm agreement between modeled nanopipette parameters and the experimental nanopipette behavior.[6]

A schematic of the FEM simulation domains for the basic surface charge model (applied to both *E. coli* and *B. subtilis*) are shown in Figure S-3. Figure S-4 shows the geometry for the more detailed gram-positive *B. subtilis* model. All boundaries that are not specifically labelled (grey) on model schematics were set as no flux boundaries with no surface charges applied. A no-slip flow condition was applied to these boundaries and the pipette walls (blue on Figure S-3 and Figure S-4). The remaining boundary conditions are described in Tables S-5 and S-6. The nanopipette potential, $V_{Tip}$, was applied to the upper boundary within the nanopipette, labelled B1, located 1 mm above the tip of the nanopipette. Boundary (B2), at a sufficient 1 mm from the nanopipette opening, was held as ground ($V$ = 0 V).

##### SI-6.1 Simple FEM model for bacterial charge

A schematic depicting the simple FEM model with a planar representation of the bacteria is shown in Figure S-3.

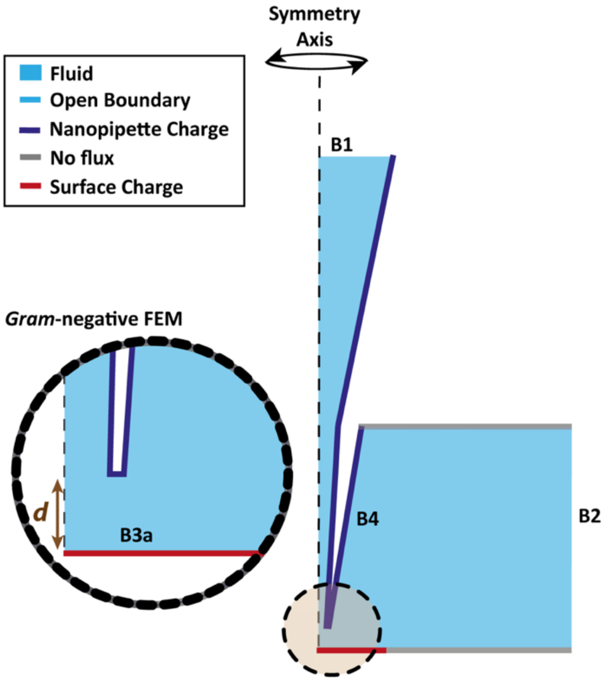


***Figure S-3:*** *Schematic (not to scale) of FEM simulation domain for the basic model.*

In the simpler, planar model the sample surface is treated as an impermeable charged insulator. Therefore, the plane labelled "B3a" (red) in Figure S3, with a radial length of 5 µm, is set to a no-flux condition and the surface is modelled to hold a (variable) charge. The analogous problem has been treated for a large planar surface in a number of our previous papers,[7,8] and this model gave identical results, and so could be applied to both the bacteria and background substrate. Boundary conditions are summarized in Table S-5. As shown B1 was set to match $V_{Tip}$, which in the case of the scanned-potential SICM simulations involved simulating a time-dependent potential sweep, such as done experimentally (50 mV to -500 mV, then reversed to 500 mV, then returned to -50 mV, with a scan rate of 1 V/s).[8]

**Table S-5:** Boundary conditions implemented in the insulating FEM simulations

| **Boundary (as shown in Figure S3)** | **Concentration/Flux condition** | **Potential/charge condition** |
|---|---|---|
| B1 | $C_i$ = 50 mM | $V = V_{Tip}$ |
| B2 | $C_i$ = 50 mM | $V$ = 0 V |
| B3a | $J_i$ = 0 | $\sigma$ = varying (-150 to 50 mC $m^{-2}$) |
| B4 | $J_i$ = 0 | $\sigma$ = -30 mC $m^{-2}$ |

The system describing the insulating simulations was defined using the following differential equations. The flux $J_i$ of each chemical species $i$, was described by the Nernst-Planck equation:

$$J_i = -D_i \nabla c_i - z_i \frac{F}{RT} D_i c_i \nabla \phi + c_i u \quad (1)$$

To solve the Nernst-Planck equation, we model the electric potential, $\phi$, using the Poisson equation:

$$\nabla^2 \phi = -\frac{F}{\varepsilon \varepsilon_0} \sum_i z_i c_i \quad (2)$$

In Equations 1 and 2, $D_i$, $z_i$ and $c_i$ are, respectively, the diffusion coefficient, charge number, and concentration of a given chemical species, $i$. $F$, $R$ and $T$ are the Faraday constant, gas constant and absolute temperature (298 K) respectively, $\varepsilon$ is the dielectric permittivity of the solution (set as 78 F $m^{-1}$), and $\varepsilon_0$ ($8.85 \times 10^{-12}$ F $m^{-1}$) describes the vacuum permittivity. The diffusion coefficients at infinite dilution for $K^+$ ($1.96 \times 10^{-5}$ $cm^2$ $s^{-1}$) and $Cl^-$ ($2.05 \times 10^{-5}$ $cm^2$ $s^{-1}$) were taken from the CRC handbook.[9] These values are reasonable because of the sufficiently dilute concentrations and the self-referencing nature of experiments.[10] The fluid velocity, $u$, is due to electroosmotic flow. This was described by the incompressible Navier-Stokes equation with electroosmotic flow (EOF) incorporated (Equation 3), where $p$ is the pressure, $\rho$ is the density of solution (set to 1000 kg $m^{-3}$), and $\mu$ is the solution viscosity (set to 0.001 Pa $s^{-1}$):

$$u \nabla u = \frac{1}{\rho}\left(-\nabla p + \mu \nabla^2 u - F\left(\sum_i z_i c_i\right) \nabla \phi\right) \quad (3)$$

##### SI-6.2 Extended FEM model for gram-positive bacteria

As shown in the FEM simulation depicted in Figure S-4, the bacterium adhered to the surface is represented as a hemisphere with a radius of 500 nm, similar to that found in TEM images (Figure S-7). A spherical shape was used as even though the charge values between geometries under standard modelling was found to be very similar, it was of interest to see the effect of cell wall thickness on simulated charge values, which could be more accurately represented using the spherical model.

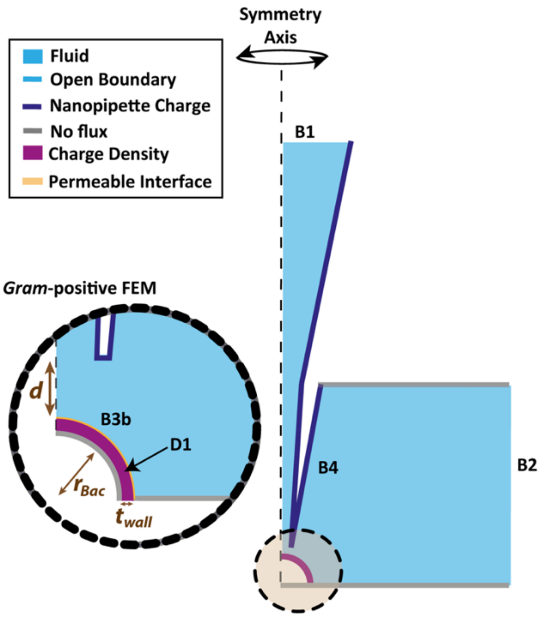


***Figure S-4:*** *Schematic (not to scale) of FEM simulation domain for the extended model of the gram-positive bacteria.*

Table S-6 demonstrates the boundary/domain conditions applied in the extended gram-positive FEM simulations. Instead of an insulating charged surface, the sample surface is defined as a permeable domain (labelled D1 on Figure S-4 - purple) with a set concentration of fixed stationary negative charge ($\rho_f/F$). The thickness of the peptidoglycan cell wall ($t_{wall}$) was estimated as 34 nm from cryo-TEM images (Section SI-7) and literature values.[11] Subsequently, a range of values from 30-70 nm were explored to examine the effects of the thickness of the cell wall on FEM simulation results.

**Table S-6:** Boundary/domain conditions implemented in the custom gram-positive FEM simulations.

| **Boundary (as labelled on Figure S4)** | **Concentration/Flux condition** | **Potential/charge condition** |
|---|---|---|
| B1 | $C_i$ = 50 mM | $V = V_{Tip}$ |
| B2 | $C_i$ = 50 mM | $V$ = 0 V |
| B3b | $J_{Cl}$ = 0 | N/A |
| B4 | $J_i$ = 0 | $\sigma$ = -30 mC m$^{-2}$ |
| D1 | $C_{Cl}$ = 0 mM | $\rho_f/F$ = *varying (0 to 500 mM)* |

An uncharged boundary is considered beneath the D1 domain, and a permeable interface above (labelled as B3b in Figure S-4, shown in yellow). The permeability of negative ions at B3b was varied in order to assess the effect of anion exclusion from the cell wall. The inclusion of a domain representing a periplasmic space with the same physical properties as the bulk solution was also examined. This was introduced as an additional domain beneath the cell wall domain (replacing the grey no-flux boundary) and had variable electrolyte concentration within.

As before, the flux $J_i$ of each chemical species $i$, was described by the Nernst-Planck equation (Equation 1). In this case, however, the Poisson equation (Equation 2) is adjusted to account for the incorporated stationary fixed charges ($\rho_f$) within the D1. Thus, the electric potential $\phi$ is given by;

$$\nabla^2\phi = -\frac{1}{\varepsilon\varepsilon_0}(\rho_f + F(\textstyle\sum_i z_i c_i)) \quad (4)$$

This new model essentially treats the cell wall as a soft polyelectrolyte layer, and has previously been used to calculate the Donnan potential within the cell wall for various species.[12] It is also similar to previous considerations of interfaces in electrophoretic measurements.[13,14] To estimate $\rho_f$, we considered the charge density of the *B. subtilis* cell wall.

This has been calculated previously using electrokinetic theory, providing charge concentration values ($\rho_f/F$) of approximately 15-25 mM.[15] However these measurements were found to be highly dependent upon strain, electrolyte composition, substrate, and pH.[14]

A variety of model variables were explored in this work to determine their effect of calculated normalized currents across a range of $\rho_f/F$ values, extending up to 500 mM. These varied parameters have been summarized in Table S-7, where the geometry parameters (labeled in brown in Figure S-3 and S-4) are separated from the non-geometry parameters within the model. With the cell wall domain (D1), the relative dielectric permittivity, $\varepsilon$, was varied at values of 7, 20 and 78 (values taken from literature from electrostatic force microscopy measurements),[16] but was found to have no significant effect upon the approach or normalized currents (shown in SI-11).

Within the cell wall domain (D1) the effective diffusion coefficients ($D_{eff}$) were varied to assess the effect of the tortuosity of the cell wall. Alteration of the diffusion coefficient in polyelectrolyte layers is due to the tortuosity of the channels which form the free space of the layer and interaction with the cell walls. $D_{eff}$ was calculated as a product of the diffusion coefficient in ideal solutions ($D$) and the relative mobility within the wall ($\mu_{wall}$) compared to solution. This factor can be estimated by the Renkin equation,[17] based upon the pore size of the *B. subtilis* cell wall (2.12 nm),[18] and the hydrodynamic radius of ions.[19] This gives estimates for $\mu_{wall}$ in the range of 0.5 to 0.75, however a wider range of 0 to 1 was explored in simulations. The effect of varying parameter values is shown in more detail in SI-11.

**Table S-7:** Parameters used for the simulations, relating to the geometry and in other parts of the simulations.

| **Geometry parameters** | | |
|---|---|---|
| ***Symbol*** | ***Value*** | ***Description*** |
| $d$ | Varied, up to 3 μm | Nanopipette-substrate separation distance, either approach distance (typically in the order of 20 nm, depending on nanopipette) or bulk (3 μm) position |
| $r_{Bac}$ | 500 nm | Radius of the spherical bacteria substrate |
| $t_{wall}$ | 30 to 70 nm | The thickness of D1, representing the varying cell wall thickness |
| **Non-geometry parameters** | | |
| ***Symbol*** | ***Value*** | ***Description*** |
| $C_i$ | -50 mM | Initial concentration of KCl in the bulk and nanopipette |
| $\rho_f/F$ | Varied, 0-500 mM | Charge concentration applied to the gram-positive cell wall for extended model |
| $z_i$ | $K^+$: 1, $Cl^-$: -1 | Charge of ions in solution ($K^+$ and $Cl^-$) |
| $D_i$ | $K^+$: 1.96 × $10^{-5}$ $cm^2$ $s^{-1}$, $Cl^-$: 2.05 × $10^{-5}$ $cm^2$ $s^{-1}$ | Diffusion coefficients of species |
| $\mu_{wall}$ | Varied, 0 to 1 | A factor for ion mobility in the cell wall domain (D1), affecting the calculated diffusion coefficient in the cell wall ($D_{eff}$) |
| $\varepsilon$ | 7, 20 or 78 | Dielectric permittivity of the cell wall (D1) |

##### SI-6.3 FEM Simulation framework

First, the experimental working approach distance between the nanopipette and the surface was estimated by steady-state simulations with the boundary at the top of the nanopipette (Boundary B1, Figure S-3 and S-4) held at $V_a$ (Figure 2, main manuscript). Then, the approach distances were taken from the simulated nanopipette-substrate separation at which the ionic current normalized to the bulk current reached the experimental approach set point of a 2 % decrease from bulk current. Time-dependent simulations of the 20 ms potential pulse were performed by stepping the voltage to $V_p$ (Figure 2, main manuscript), using the corresponding steady-state approach simulation as starting conditions (see above). The time-dependent simulation step was run for both a large nanopipette-substrate separation, *i.e.* bulk conditions, and for the working approach distance calculated from the previously steady state simulation. Simulated currents at working distance were then normalized by the ones simulated at bulk conditions resulting in normalized current values. This framework was performed for all simulation regardless to which model (planar or spherical) was used.

#### SI-7 Electron microscopy analysis of bacteria

Bacteria cultures grown overnight from freezer stocks in M9m media were added to carbon-laced copper or Formar/Carbon on 200 Mesh copper TEM grids, and then imaged using a Zeiss Supra 55VP, or a JEOL 2100 TEM. An SEM image of *E. coli* is shown in Figure S-5, where the dimensions are consistent with those found in the SICM topography map in Figure 3 (width of approximately 0.8 μm), SEM images of the *B. subtilis* (Figure S-6 and S-7) show an additional material surrounding the bacteria that is distinct from the TEM grids, we believe this could be attributed to extracellular polymeric substance (EPS).

Cryo-EM (JEOL 2100 TEM) was also employed to observe the structure of the *B. subtilis* cell wall (Figure S-8). Samples were frozen in liquid ethane using a Leica EM GP2 cryo plunge freezer. Multiple cells from the same sample were imaged, and cell wall measurements were taken at 3 different locations of each cell to calculate the average thickness of each cell wall fraction, see Table S-8. These mean values were used for the FEM simulations.

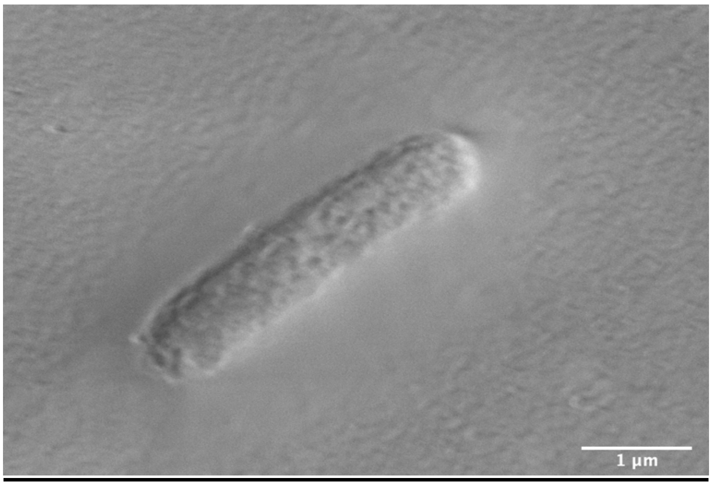


***Figure S-5:*** *Scanning Electron Microscopy (SEM) image of Escherichia coli (wild type) cell, observed on carbon-laced copper TEM grids, electron high tension (EHT) = 0.500 kV.*

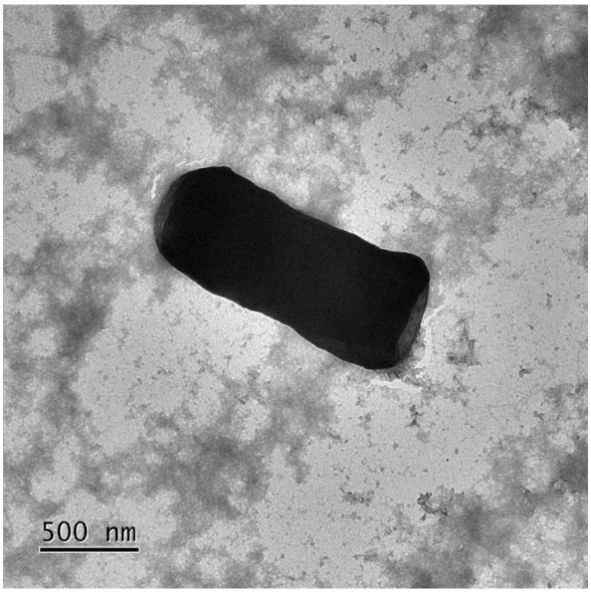


***Figure S-6:*** *Scanning Electron Microscopy (SEM) image of Bacillus subtilis (Δhag) cell, observed on a Formar/Carbon copper TEM grid. EHT = 200 kV.*

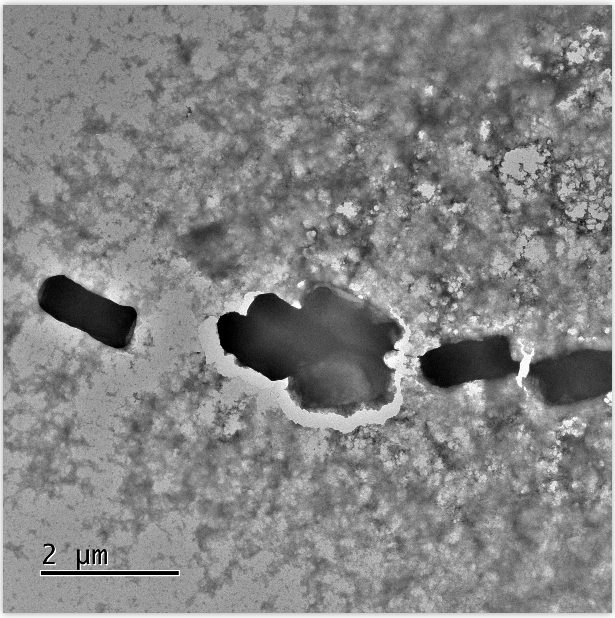


***Figure S-7:*** *Scanning Electron Microscopy (SEM) image of Bacillus subtilis (Δhag) cell, on a carbon-laced copper TEM grids. EHT = 200 kV.*

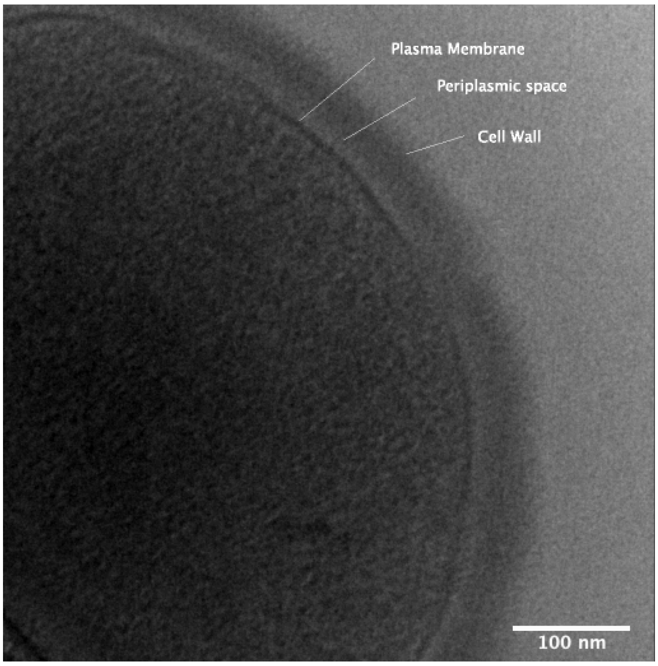


***Figure S-8:*** *Cryo-EM image of Bacillus subtilis (Δhag) cell, observed on carbon-laced copper TEM grids. EHT = 200 kV.*

**Table S-8:** Cell wall thickness measurements from cryo-EM images of Bacillus subtilis (Δhag) cells (*n*=12), measured at 3 distinct locations along the cell wall.

| | **Range (nm)** | **Mean (nm)** | **STD (nm)** |
|---|---|---|---|
| Total Cell Wall | 51-61 | 54.8 | 3.2 |
| Cell Wall (Peptidoglycan layer) | 31-40 | 34.0 | 3.2 |
| Periplasmic space | 12-16 | 14.6 | 1.4 |
| Periplasmic membrane | 7-10 | 8.3 | 1.2 |

#### SI-8 Further analysis of pulse-potential *E. coli* scan

As discussed in SI-6 and in the main manuscript, to explore the effects of the spherical topography of bacteria on the model results, in a 2D axisymmetric cylindrical geometry, FEM simulations compared treating the bacteria as a planar substrate compared to a spherical substrate (radius of 0.5 μm).

Figure S-9A shows the charge from the *E. coli* pulse potential scan using the spherical topography model instead of the planar topography model, as discussed in Figure 3 (main text). There is, at most, a difference of 10 mC m$^{-2}$ between the two models, with more positive charge values calculated by the spherical model. Figure S-9B shows the approach curve for the spherical model, used for calculating the experimental approach height (see Section SI-6.3), where the 2 % threshold is met at similar heights to that expected with the planar model (30 nm).

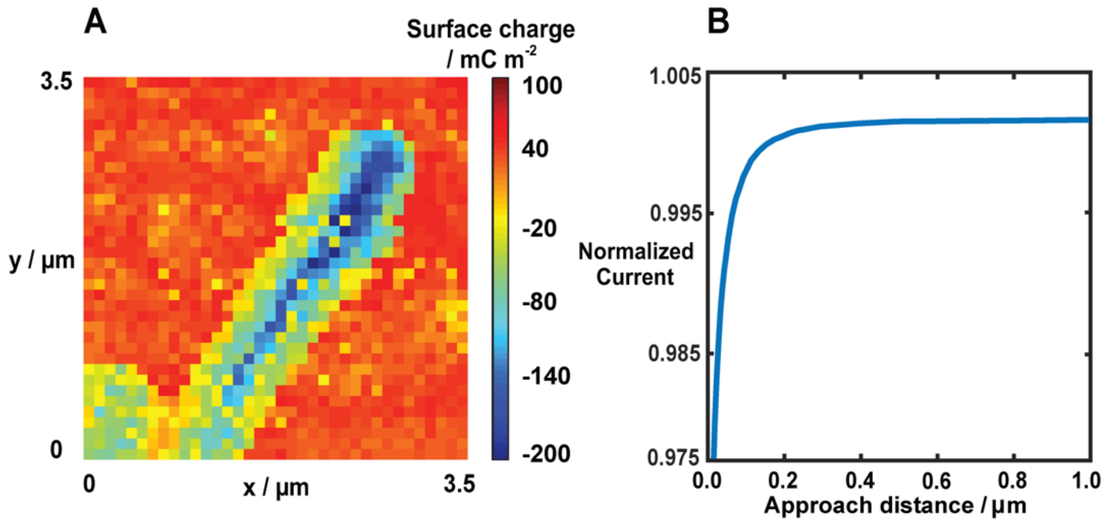


***Figure S-9.*** *(A) Charge values (mC m*$^{-2}$*) for E. coli pulse potential scan (Figure 3, main manuscript) as calculated using the spherical FEM simulations, (B) Simulated approach curve using the spherical FEM.*

We also considered whether the pressure gating on ion channels could be activated by the pressure generated by the movement of the SICM tip. Figure S-10 shows how the pressure (mbar) changes down a vertical line along the axis of symmetry of nanopipette, starting from 100 nm into the nanopipette end, and extending to 500 nm outside of the nanopipette. The pressure drops considerably within the first 50 nm exiting the nanopipette, and within 300 nm the pressure is effectively 0 mbar. Three times points across the -500 mV SICM pulse (0, 3 and 15 ms) are shown, demonstrating that the pressure is irrespective of the time during the pulse.

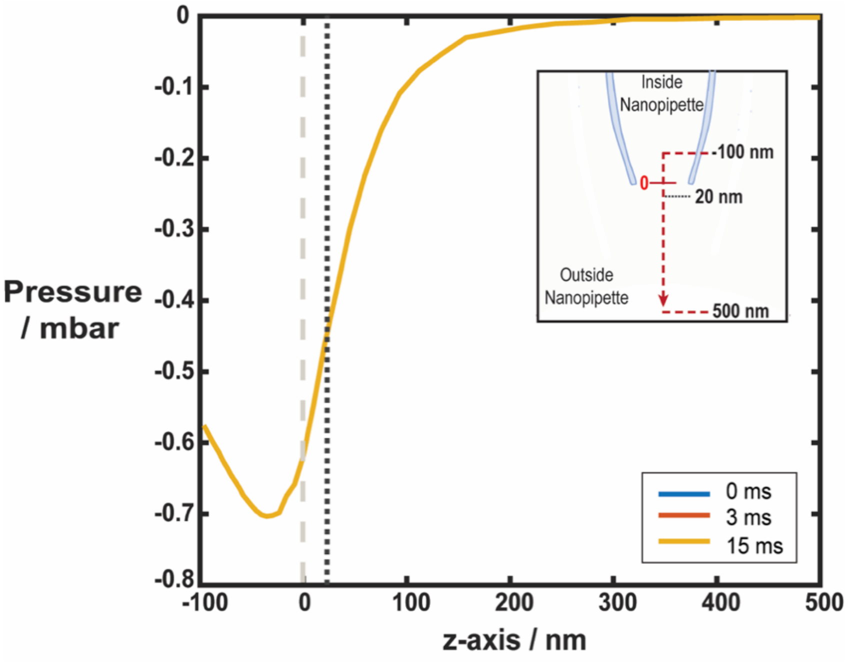


***Figure S-10.*** *Pressure profile vertically intersecting the nanopipette from 100 nm into the nanopipette end, to 500 nm outside of the nanopipette. The nanopipette opening at the grey dashed line, where the estimated nanopipette-substrate separation (20 nm) used for surface measurements is shown in black dotted line. Three time points across the pulse ($V_p$ = -500 mV) are shown overlaid, at the beginning of the pulse (0 ms), during the ramping of current into steady state (3 ms), and at an established steady-state current time (15 ms).*

Taking values from Figure S-10, at the estimated approach height above the cell, 20 nm, the pressure will be approximately -0.45 mbar (-45 Pa). Using values from Martinac *et al.* (1987) patch-clamp experiments, the pressure required for probable opening (above 60 %) of the ion channels in *E. coli*, with approximately -20 mV being applied from the nanopipette during pulse, is *ca.* -26.6 mbar (-2.66 kPa).[20] Therefore, even under a potential the -20 mV modelled at the surface, the required pressure needed to open channels within *E. coli* is 50-fold greater than is achieved in the pulse experiments. Whilst these values are based on *E. coli* from patch-clamp experiments, there are similarities in the families of mechanosensitive channels across gram-positive and gram-negative bacteria.[21]

#### SI-9 FEM simulations for the scanned-potential *E. coli* scan

The scanned-potential *E. coli* map was calibrated using a series of *i-V* curves with matching conditions to the SICM experiment (+/- 500 mV, 1 V/s), where the model was executed at the approach height under a series of surface charges applied to the substrate below (-150 to 100 mC $m^{-2}$). The currents at extreme potentials (+/- 500 mV) near the surface were normalized against bulk values forming two calibration curves shown in Figure S-11B. The normalized current and calibrated charge at the nanopipette potential of 500 mV in the scan is shown in Figure S-11C/D, with the equivalent at -500 mV at the nanopipette shown with no background in Figure 5 (main manuscript), and the raw data included the heterogenous background in shown in the Figure S-11E/F. The charge values across the bacteria are similar between both extremes, with cell surface values of approximately -70 to -120 mC $m^{-2}$, demonstrating how this regime can be used to calibrate the images reliably in both potential directions.

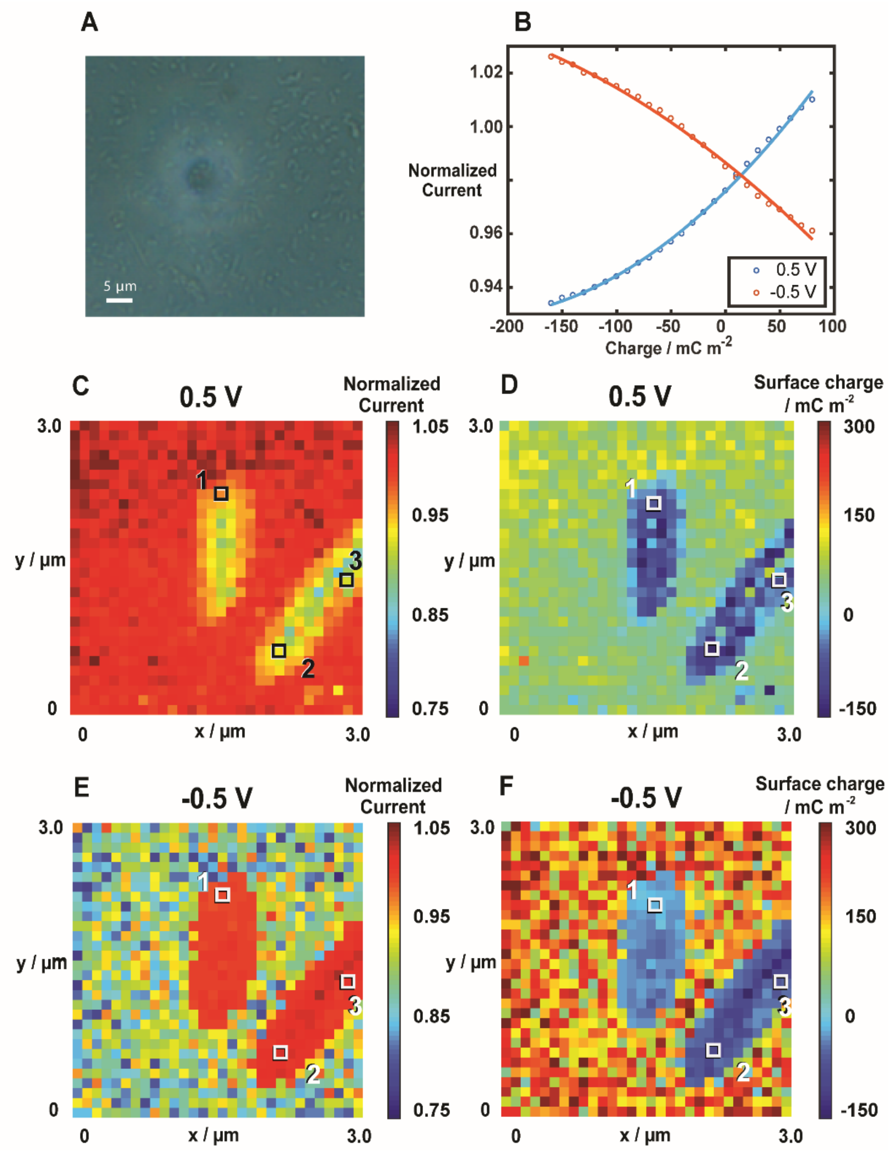


***Figure S-11.*** *(A) Optical micrograph of the bacteria before scanning, scanned area is within the halo caused by the optical properties of the nanopipette (B) Simulated calibration curve for the E. coli scanned-potential scan described in Figure 5 (main text), shows calibration plots taken from data plots at -500 and 500 mV at the nanopipette on the i-V scans with different charges applied to the bacterial substrate. (C) Normalized current and (D) calibrated charge of the scanned-potential scan (from Figure 5) at 500 mV applied to the nanopipette, with the equivalent full data for the -500 mV pulse shown in (E) and (F).*

It is noted that the charge values across the background Cell-Tak substrate in both potential directions are positive, but with higher values in the negative swing. Importantly, over the bacteria the calculated charges are similar in both potential directions, which supports our argument that the gram-negative bacteria act as an insulating substrate.

#### S-10 SICM of *B. subtilis* in biological media

In addition to the conditions reported in the paper, bacteria were studied in the physiological growth medium 9m (described in SI-2) as the bulk bath and nanopipette solution. These additional scans were used to determine if the relatively high values of normalized currents found in *B. subtilis* related to osmotic stress on the bacteria due to the 50 mM KCl. Figure S-12 shows a SICM scan of the *B. subtilis* Δ*hag* using M9m media.

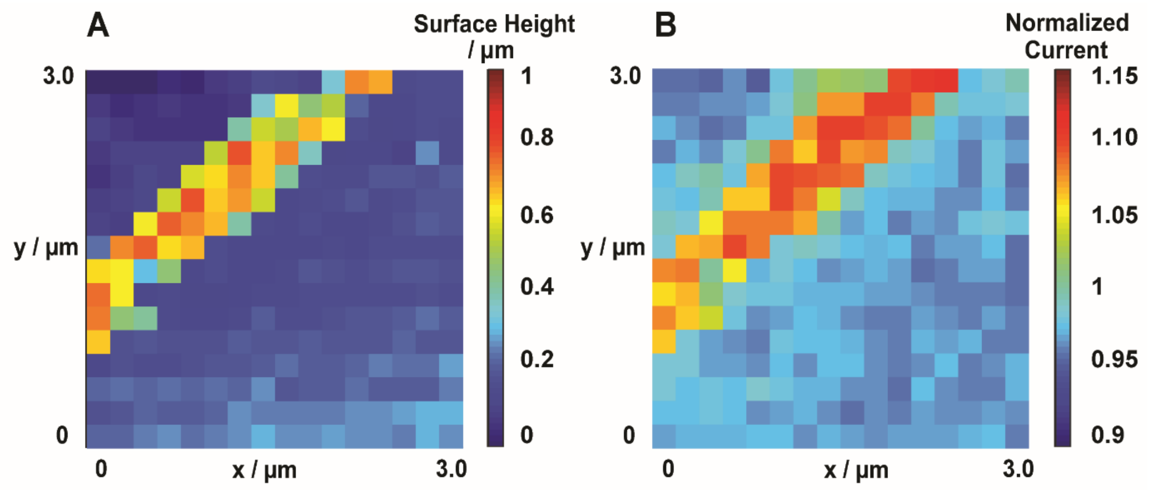


***Figure S-12.*** *SICM topography (A) and normalized current, taken from the end of the 20 ms pulse of -500 mV (B) of B. subtilis Δhag adhered to glass by Cell-Tak, using M9m (pH 7) as the nanopipette and bath solution. 50 mV approach bias utilizing a 2 % current feedback threshold. 150 nm nanopipette with 0.2 µm hopping distance*

The normalized currents across the bacteria shown in Figure S-12B are in the range of 1.08-1.12, proximal to the values observed with the *B. subtilis* scanned in 50 mM KCl (shown in Figure 6, in the main text). The slightly lower normalized current values may relate to the higher ionic concentration of M9m media, compared to 50 mM KCl. Full media speciation of the M9m using the MINEQL software produced an ionic strength concentration of approximately 240 mM. As shown previously, higher ionic strength (above 100 mM) of the electrolyte results in compression of the Debye lengths on the charged surface.[10] This means that at the comparative nanopipette height above the surface at the 2 % current threshold, there will be a greatly reduced sensitivity to surface charge with the higher strength M9m media scanning, resulting in smaller normalized current values.

#### SI-11 Further results from the extended FEM model

For *B. subtilis*, the parameters varied for exploration were $\rho_f/F$, $\mu_{wall}$, $\varepsilon$, and membrane thickness. Figure S-13A shows the approach curve simulations for *B. subtilis*, with a target approach height of 20 nm, which is independent of $\rho_f/F$ (volumetric charge concentration) as expected for the small applied approach bias of 50 mV.[22] Figure S-13B shows the effect of the peptidoglycan cell wall thickness on the normalized current across a range of wall charges. With increasing thickness of cell wall, normalized current increases.

Figure S-13C shows the small difference in calculated normalized current made by changing the dielectric permittivity within the cell wall; a permittivity constant of 20 was used in subsequent simulations, as determined for other gram-positive strains.[16] Simulated current-time responses, upon stepping the potential from 50 mV to -500 mV, are shown in Figure S-13D, with the nanopipette near the surface (distance of 20 nm) and in bulk position, i.e., 2 μm above the cell wall, with a charge density of 100 mM.

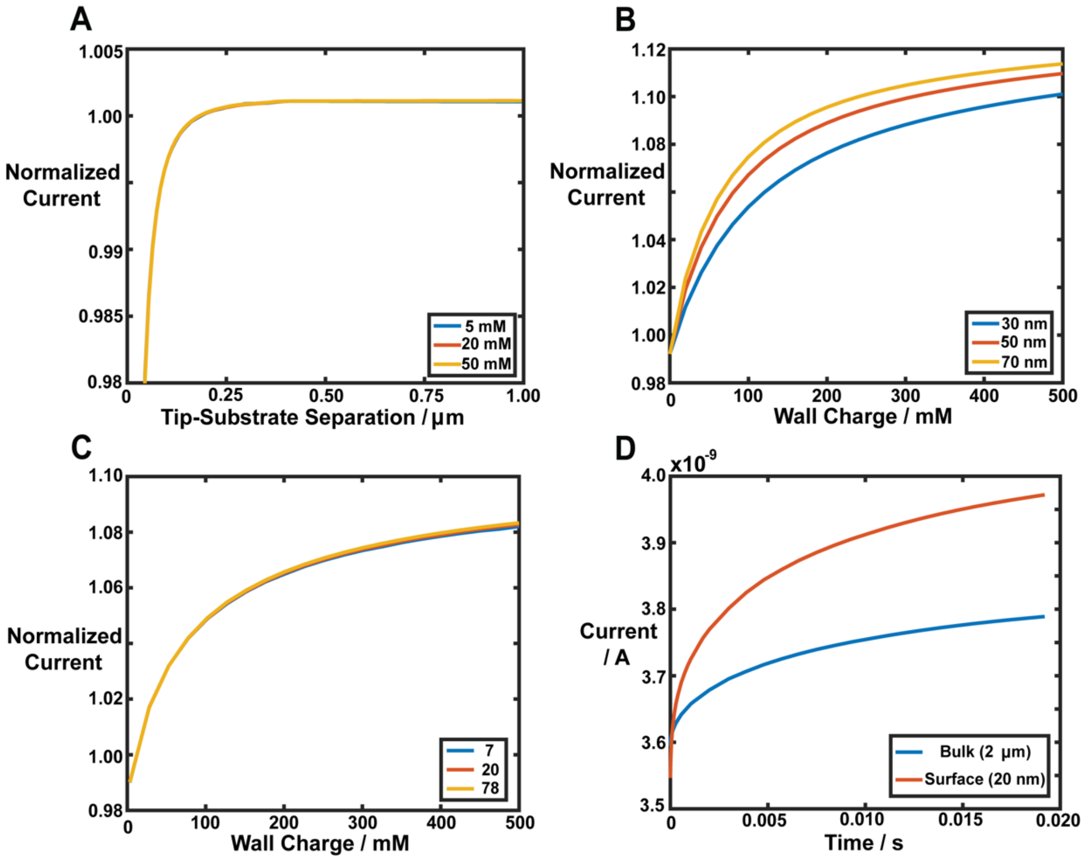


***Figure S-13:*** *FEM simulations for functional mapping of B. subtilis. (A) Simulated approaches ($V_{TIP}$ = 50 mV) to a gram-positive cell wall with several different charge density,* $\boldsymbol{\rho_f / F}$*, values defined in the key. (B) Simulated normalized current ($V_{TIP}$ = -500 mV) for different wall charge density values,* $\boldsymbol{\rho_f / F}$*, defined in the key, as a function of cell wall thicknesses, with the tip at 20 nm from the cell wall. (C) Simulated normalized current as a function of wall charge density,* $\boldsymbol{\rho_f / F}$*, values with different values of dielectric permittivity,* $\boldsymbol{\varepsilon}$*, of the cell wall (key). Nanopipette-substrate separation of 20 nm, cell wall thickness of 70 nm. (D) Example simulated current time curves for a tip potential pulse of -500 mV (from an initial value of 50 mV) near the cell wall surface (tip-surface separation of 20 nm), with* $\boldsymbol{\rho_f / F}$ *= 100 mM compared to the same pulse conditions in the bulk solution (2 µm above the surface).*

The exterior of the cell (boundary B3b on Figure S-4) was set to be impermeable to $Cl^-$ and $\mu_{wall}$ was only applied to $K^+$. This is considered reasonable as anions have been found to be excluded from the cell wall at these electrolyte concentrations.[12]

It is worth noting that a few pixels in the *B. subtilis* charge map displayed in Figure 7 (main text) show a normalized current of 1.15, and the maximum value achievable through simulation as shown in Figure 8B (main text) is approximately 1.1. Using Figure S-13D, a normalized current of 0.05 relates to approximately 200 pA of current. This 200 pA current would equate to approximately $1.9 \times 10^7$ monovalent ions (with the mobility of $K^+$) for the pulse duration of 20 ms. It is possible that some of this current could result from inducing the transfer of ions through the peptidoglycan cell wall, but this may not realistically account for the whole of the unaccounted flux if we consider only ion channel related activity. A combination of effects from ion channels, charged proteins and membrane sugars releasing ions might be important and, as described in the main text, an interesting avenue for future SICM studies would be to use it in a more intrusive mode in order to further improve biophysical models of ion dynamics on the cell surface and in the microenvironment.

(7) Page, A.; Perry, D.; Young, P.; Mitchell, D.; Frenguelli, B. G.; Unwin, P. R. *Anal. Chem.*

**2016**, *88* (22), 10854–10859.

(8) Perry, D.; Paulose Nadappuram, B.; Momotenko, D.; Voyias, P. D.; Page, A.; Tripathi, G.; Frenguelli, B. G.; Unwin, P. R. *J. Am. Chem. Soc.* **2016**, *138* (9), 3152–3160.

(9) *CRC Handbook of Chemistry and Physics*, 96th ed.; Haynes, W. M. (Editor-in-chief), Lide, D. R., Bruno, T. G., Eds.; Taylor and Francis Group: Boca Raton, 2015.
